# An amplicon panel for high-throughput and low-cost genotyping of Yesso scallop *Mizuhopecten yessoensis*

**DOI:** 10.1101/2025.11.05.686843

**Authors:** Ben J. G. Sutherland, Chen Yin Walker, Denman Moody, Chelsea Bennett, Marissa Wright-LaGreca, Chloe Shamash-McLaughlin, Karen Leask, Daniel Roth, Megan Lebeuf, Timothy J. Green

## Abstract

The Yesso scallop *Mizuhopecten yessoensis* was imported from Japan to western Canada in the late 1980s to establish an economically viable scallop aquaculture industry. Since this time, the industry in Canada has operated with existing genetic diversity within the broodstock, which is considerably limited relative to wild populations. The sector has not been able to realise its full potential in part due to idiopathic hatchery failures and farm stock collapses due to disease outbreaks associated with the intracellular bacterial pathogen *Francisella halioticida*. To support Yesso scallop production and breeding, here we generate a low-density, genotyping-by-sequencing amplicon panel using single nucleotide polymorphism (SNP) markers that are evenly spaced across the *M. yessoensis* genome and that show high heterozygosity in Canada and Japan. The panel can also exploit the high genetic polymorphism of the *M. yessoensis* genome, with *de novo* SNP calling identifying over 2,500 high quality SNPs within the 579 sequenced amplicons. We demonstrate the utility and versatility of this new genotyping tool for breeding applications including parentage assignment, low density family-based genome-wide association study, trait heritability evaluation to determine potential for genomic selection, and species differentiation (against the weathervane scallop *Patinopecten caurinus*). We did not find any genomic regions significantly associated with *F. halioticida* resistance but did identify potential for genomic selection. We could separate the two species based on genotypes, and we did not see evidence of a past *M. yessoensis* x *P. caurinus* hybridization event within the *M. yessoensis* breeding population at Vancouver Island University. This low-cost genotyping panel is expected to accelerate selective breeding improvements for *M. yessoensis* in Canada and elsewhere.

## Introduction

The marine bivalve Yesso scallop *Mizuhopecten* (previously *Patinopecten*) *yessoensis* is endemic to Asia and is a highly valued protein source served as a delicacy by restaurants globally. It is an economically valuable species in China, Russia, Korea, and Japan, and has been introduced for culture in France, Morocco, and Canada (reviewed by Kawahara et al., 2019). In Japan, scallops have the highest productivity within sea farming industries (Kosaka, 2016). China produced 1.42 million tonnes of scallops in 2012, and although Yesso scallop production is not the species produced at the highest mass, it is the highest in commercial value (Guo and Luo, 2016). There is interest in scallop culture on the west coast of Canada, in part due to its success in Japan and strong market demand (Parsons et al., 2016). However, the current production of Yesso scallop in British Columbia (BC), Canada, is far below its potential (Holden et al., 2019). BC produced 695 tonnes of scallops in 2010, and only 96 tonnes in 2023 (DFO, 2023). There is potential for establishing productive scallop aquaculture industries globally, but significant genetic work will be needed to achieve this goal (Bourne, 2000).

Compared to the major aquaculture bivalve species Pacific oyster *Crassostrea* (*Magallana*) *gigas*, Yesso scallop has been domesticated more recently (Bourne, 2000) and poses more challenges for hatchery production (Robert and Gérard, 1999). In western Canada, breeding program and aquaculture populations have much lower effective population sizes (*N*_e_) than wild populations in Japan (Sutherland et al., 2023). This lower genetic diversity could result in inbreeding, especially when breeding program pedigrees are incomplete or if genetic similarity between founders is unknown. Inbreeding can result in poor growth or survival, limiting production. Yesso scallop was first grown commercially in Canada in the late 1980s (Bourne, 2000), but has no self-recruiting populations in the area, unlike Pacific oyster (Quayle, 1988; Guo, 2009; Sutherland et al., 2020). Self-recruiting or wild populations provide an option to replenish broodstock in the event of a loss of diversity, and therefore in Canada replenishment would require new international imports. As a result, hatcheries are left with the current broodstock supply, highlighting the value in preserving and making best use of this existing genetic diversity.

A rapid and low-cost amplicon panel for genotyping is expected to benefit Yesso scallop aquaculture in western Canada and elsewhere, similar to that developed for Pacific oyster (Sutherland et al., 2024), due to its ease of use, affordability, and reliability (Meek and Larson, 2019). This tool could be used to detect pedigree errors or contaminated families in breeding programs to protect directional breeding progress (see Hedgecock and Davis, 2007), to avoid crossing related individuals when pedigrees are unavailable or uncertain, and to enable genomic selection including with pedigree-based genome-wide imputation (Delomas et al., 2023; Sutherland et al., 2026). In fact, genomic selection has been shown to have potential in Yesso scallop from simulation studies for panels with densities from 5K to 250K single nucleotide polymorphisms (SNPs; Dou et al., 2016), but has not yet been put into application.

Amplicon panels are particularly effective at genotyping shellfish due to their high genetic polymorphism (e.g., a SNP every 25 bp in non-coding regions of Pacific oyster; Hedgecock et al., 2005; Sauvage et al., 2007). Amplifying genomic regions rather than requiring the presence of specific SNPs should increase the lifespan and scope of the platform (Thompson et al., 2025). Given the high polymorphism turnover expected due to genetic drift and sweepstakes reproductive success (Hedgecock and Pudovkin, 2011), and the emergence of new mutations via high mutation rates (Plough, 2016), a low-density panel is most effective if it can flexibly characterize any SNPs within a set of specified windows. Amplicon panels can also be used for microhaplotype genotyping, providing a more powerful marker type than single SNPs for various applications, including *N*_e_ estimation (Sutherland et al., 2023) and parentage assignment (Baetscher et al., 2018; Thompson et al., 2025).

The Yesso scallop industry in BC has been specifically challenged by diseases caused by parasites and pathogens (Gillespie et al., 2012). One such agent is *Francisella halioticida*, an intracellular bacterial pathogen originally described in giant abalone *Haliotis gigantea* in Japan (Kamaishi et al., 2010; Brevik et al., 2011). It is a significant threat to the industry in Canada. In 2015, *F. halioticida* was associated with a large-scale mortality event of juvenile Yesso scallops in BC that resulted in 40% mortality (Meyer et al., 2017). *F. halioticida* has been confirmed as causative in Yesso scallop mortalities (Kawahara et al., 2019). Infections are associated with lesions, slow growth, and mortality (Meyer and Itoh, 2025). *F. halioticida* is likely widely distributed in Japan, with prevalence increasing during the summer when high mortality occurs (Meyer and Itoh, 2025). Positive *in situ* hybridization detections of *F. halioticida* in archived tissue suggests the presence of the bacterium in BC since the late 1980s (Meyer et al., 2017). Infection susceptibility is likely related to environmental stressors compromising physical or immunological defenses (Kawahara et al., 2019), but may also be related to genetic factors. Reducing the susceptibility of hatchery-grown Yesso scallop through selective breeding (Bourne, 2000) may reduce the burden of this pathogen on the industry. Another approach to protecting Yesso scallop from disease has been to produce Yesso x weathervane hybrids (Saunders and Heath, 1994), which exhibit improved disease resilience (Bower et al., 1999). This has also been demonstrated elsewhere, with hybrid vigour being observed in growth and temperature tolerance (Xing et al., 2022). It is not clear as to what extent the BC-produced hybrid lines have spread through the broodstock used in contemporary BC aquaculture.

In the present study we use previously identified SNP markers that are polymorphic in Japan and Canada (Sutherland et al., 2023) to select a subset of 600 target regions to include in an amplicon panel. Selected regions were tiled evenly across the chromosome-level genome (Wang et al., 2017; Liu et al., 2020), with preference to target high heterozygosity markers in both Canada and Japan (Sutherland et al., 2023). Within the target regions, we use a *de novo* SNP detection approach to expand the variants genotyped by the panel beyond a single designed target marker per amplicon (Thompson et al., 2025). We then test out the panel in a variety of applications to test its general utility: (1) parentage application; (2) low-density genome-wide association study to *F. halioticida* disease resistance; (3) heritability of *F. halioticida* disease resistance and therefore potential for genomic selection; and (4) cross-species amplification for species discrimination between Yesso scallop and weathervane scallop *P. caurinus*. Based on these results, this panel is expected to form the basis for genetically informed selective breeding and broodstock management in BC, Canada.

## 2 Methods

### 2.1 Data sources and marker selection

Candidate SNPs were obtained from a double-digest restriction site-associated DNA sequencing (ddRADseq) study of Yesso scallops. The study included samples from a breeding program at Vancouver Island University (VIU), a commercial farm in BC, and wild-caught scallops from coastal Japan (see Data Availability; Sutherland et al., 2023). The present analysis used per population, per locus observed heterozygosity (*H*_OBS_) and Hardy-Weinberg equilibrium statistics for populations from Japan and Canada (VIU), as previously described (Sutherland et al., 2023). The Yesso scallop scaffold-level reference genome used for ddRADseq genotyping was obtained from NCBI (GCA_002113885.2; Wang et al., 2017), and a chromosome-level assembly for Yesso scallop was obtained from the genomic database MolluscDB (Liu et al., 2020) on June 19^th^, 2023 (md5: a479*2253).

Several approaches were taken to optimize marker selection: (1) select one SNP per window based on highest *H*_OBS_ in a non-overlapping tiling window approach with window size 1.7 Mbp; (2) select three SNPs per 5.0 Mbp window; or (3) select 12 SNPs per 20.1 Mbp window. When constructing windows, all chromosomes were connected in a continuous sequence, ignoring chromosome separations. Marker *H*_OBS_ used in the selection were based on the sum of the *H*_OBS_ calculated individually for populations from VIU and Japan. The best approach was determined by the average *H*_OBS_ of selected markers and the number of total markers retained.

Positions of the SNPs from the ddRADseq study were transferred from the scaffold assembly to the chromosome assembly using SNPLift (Normandeau et al., 2023) as per software defaults but with the *CORRECT_ID* flag turned off to retain the names from the original genome to match with original per locus statistics from Sutherland et al. (2023). Markers were considered for inclusion in the panel if they had *H*_OBS_ ≤ 0.5 in each of the two populations and did not significantly deviate from Hardy-Weinberg proportions (p > 0.05). Following the optimal window selection approach, additional high heterozygosity SNPs were added until the total number of selected SNPs reached 600 SNPs. The chromosome assembly contains a scaffold constructed of remaining contigs that had not yet been fit into chromosomes, named chr00 (Yuli Li, *pers. comm.*), and this scaffold was also included in marker selection and design. Additional details are provided in the *amplitools* pipeline, panel designer section (see *Data Availability*).

Two custom variants were added to the candidate markers for inclusion in the panel based on putative functional variation in the literature. These included two SNPs associated with colour variation in Yesso scallop (Zhao et al., 2017): SNP 6363 and SNP 259 at positions 22,014,997 and 25,302,657 on chromosome 11. For these two custom variants, a 401 bp window (200 bp either side of the variant) was extracted from the chromosome-level assembly, then NCBI BLAST was used to query the scaffold-level assembly on which the panel would be designed.

### 2.2 Panel design

The VCF file from Sutherland et al. (2023) was subset to only include the selected SNPs, and a bed file was created specifying 200 bp up- and down-stream from the target site using custom code (see *Data Availability*, *ms_scallop_panel*). The bed file was used with bedtools (Quinlan and Hall, 2010) to extract the specified windows from the reference genome, as well as the expected reference and alternate alleles, to prepare a design submission file (see Additional File S1). The VCF file used was produced as an output of Stacks2 (Rochette et al., 2019), a program which calls the reference allele based on the most frequently viewed variant. As such, for the panel design, the reference genome was used to update the designated reference allele based on the nucleotide present in the reference genome (see *Data Availability*, *ms_scallop_panel*). The final primer panel design was conducted by Thermo Fisher Scientific using the AgriSeq workflow, following standard manufacturer protocols.

### 2.3 Pilot study experimental designs, *F. halioticida* infection trial, and tissue collection

To evaluate the functionality of the amplicon panel, pilot studies were conducted comprising four analyses: (1) parentage within the VIU breeding program; (2) low-resolution family-based genome-wide association study (GWAS) for resistance against *F. halioticida*; (3) heritability evaluation for resistance to *F. halioticida*; (4) cross-species genotyping and species delineation between weathervane scallop *P. caurinus* and Yesso scallop. Additional sampling tests were conducted to evaluate optimal DNA sampling methods. Table 1 and the text below provides a description of these experiments and analyses.

**Table 1.**
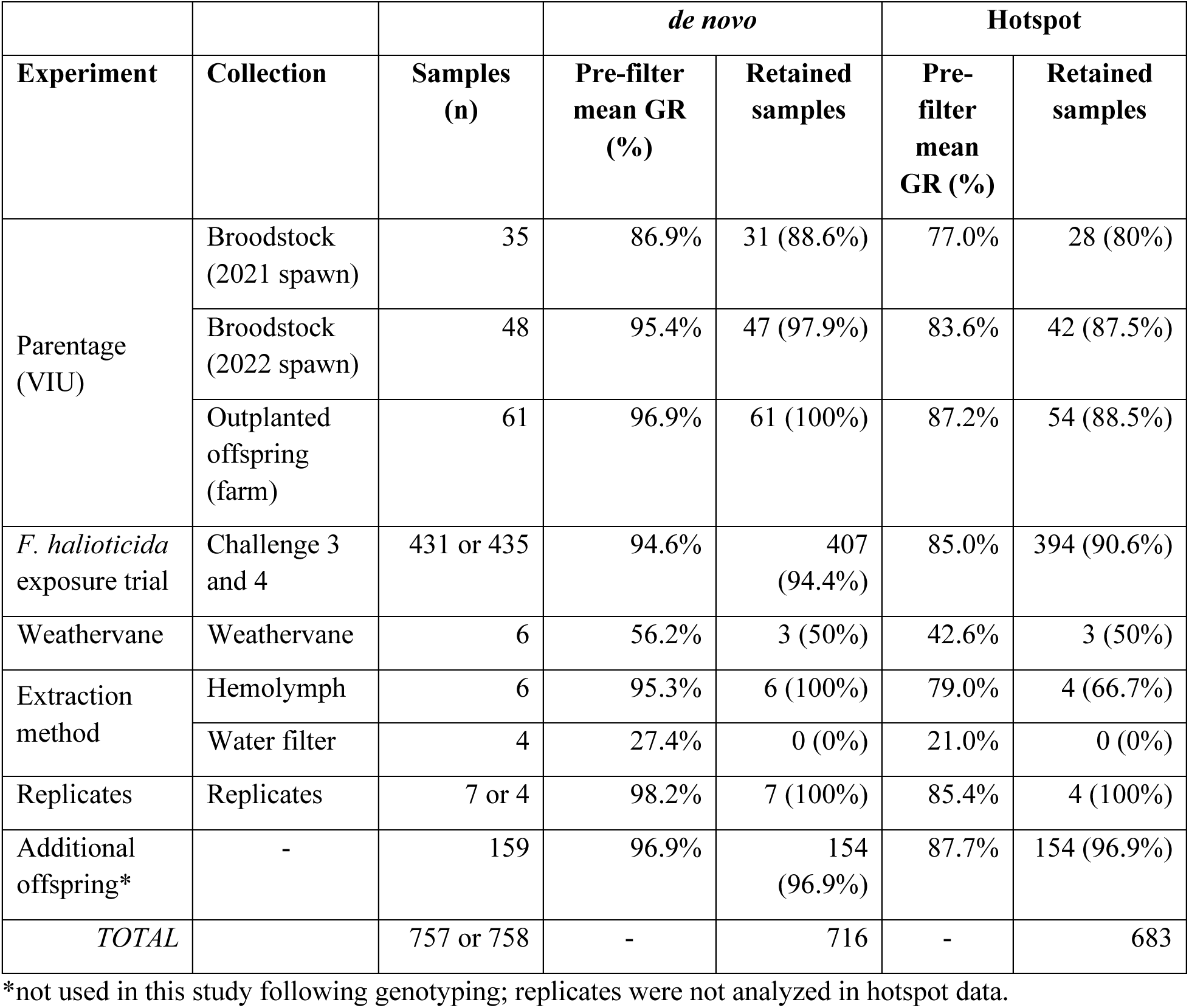
The pilot project was comprised of a parental study, an analysis of genetic basis of resistance to *F. halioticida* within families in the VIU breeding program, and a cross-species amplification of weathervane scallop alongside the Yesso scallops. Genotyping was conducted with all samples together and included 2,592 filtered SNP variants in the *de novo* dataset, and 581 SNPs in the hotspot dataset. Samples were individually filtered based on genotyping rate (GR), and the number of samples passing filters are shown (‘Retained samples’) as a percentage of the total input samples. When the input number of samples column has two values for a row, the first is for the *de novo* and second for the hotspot, as in some cases there was no sequence data generated.

|  |  |  | *de novo* |  | Hotspot |  |
| --- | --- | --- | --- | --- | --- | --- |
| Experiment | Collection | Samples (n) | Pre-filter mean GR (%) | Retained samples | Pre-filter mean GR (%) | Retained samples |
| Parentage (VIU) | Broodstock (2021 spawn) | 35 | 86.9% | 31 (88.6%) | 77.0% | 28 (80%) |
|  | Broodstock (2022 spawn) | 48 | 95.4% | 47 (97.9%) | 83.6% | 42 (87.5%) |
|  | Outplanted offspring (farm) | 61 | 96.9% | 61 (100%) | 87.2% | 54 (88.5%) |
| *F. halioticida* exposure trial | Challenge 3 and 4 | 431 or 435 | 94.6% | 407 (94.4%) | 85.0% | 394 (90.6%) |
| Weathervane | Weathervane | 6 | 56.2% | 3 (50%) | 42.6% | 3 (50%) |
| Extraction method | Hemolymph | 6 | 95.3% | 6 (100%) | 79.0% | 4 (66.7%) |
|  | Water filter | 4 | 27.4% | 0 (0%) | 21.0% | 0 (0%) |
| Replicates | Replicates | 7 or 4 | 98.2% | 7 (100%) | 85.4% | 4 (100%) |
| Additional offspring* | - | 159 | 96.9% | 154 (96.9%) | 87.7% | 154 (96.9%) |
| *TOTAL* |  | 757 or 758 | - | 716 | - | 683 |

\*not used in this study following genotyping; replicates were not analyzed in hotspot data.

**Table 2.** Fixed differences in loci genotyped in the Yesso scallop vs. weathervane scallop analysis, including Yesso scallops genotyped by ddRADseq (denoted by asterisk) from a BC farm, Japan (JPN) wild, or the VIU breeding program, or by amplicon panel (VIU breeding program).

|  | Yesso (BC)* | Yesso (JPN)* | Yesso (VIU) | Weathervane |
| --- | --- | --- | --- | --- |
| Yesso (JPN)* | 0 |  |  |  |
| Yesso (VIU) | 0 | 0 |  |  |
| Weathervane | 20 | 17 | 16 |  |
| Yesso (VIU)* | 0 | 0 | 0 | 17 |

The parentage study included broodstock from spawn events in 2021 (scallop family run 2; SFR2) and 2022 (SFR3 and SFR4). Tissues were non-lethally sampled from broodstock by cutting a small section of mantle and freezing in a -80°C portable freezer and from offspring using a syringe to extract a small volume of hemolymph (< 100 µl), spinning down the volume in a centrifuge, then flash freezing the cell pellet on dry ice. Broodstock were sampled at the Deep Bay Marine Field Station of VIU, and outplanted offspring were collected from a farm in British Columbia (BC) following one year in the field. Genomic DNA was extracted and purified using the Monarch Spin Genomic DNA Purification kit (New England Biolabs) following the tissue protocol for broodstock and the blood protocol for offspring. The true parents of the outplanted (farm) offspring were the broodstock used in the 2022 spawn events (i.e., SFR3 and SFR4). An additional set of offspring were also genotyped from the 2021 (SFR2) spawn event but were not used for any analyses here as they are part of a different project.

*F. halioticida* challenges were conducted on scallops produced by the 2021 (SFR2) spawn event using an isolate gifted by G. Meyer, with pathogenicity in *M. yessoensis* previously determined in disease studies (Meyer et al., 2017; Kawahara et al., 2019). Disease challenge protocols were adopted from Kawahara et al. (2019), and *F. halioticida* was grown on modified Eugon agar (Kawahara et al., 2018). In brief, the infection trial was conducted by injecting 50 µL of *F. halioticida* inoculum (OD_600_=0.05) into the adductor muscle of each scallop (shell height: 36.3 ± 8.3 mm) using a 26-gauge needle attached to a multi-dispensing pipette. The scallops were housed in aerated 10 L aquaria filled with 1 μm-filtered seawater maintained at 16°C, with morbidity assessed daily. Morbidity was measured for five days, after which time the disease challenge ended. Scallops were considered moribund or deceased when their shell valves were gaping and failed to close after gently touching the scallop with a gloved finger. Upon morbidity or the end of the five-day trial, each individual scallop was measured for size, and recorded based on whether the scallop had died during the trial, and if so, the day of death. A control treatment containing a random subset of scallops (n = 18) that were not injected with *F. halioticida* but instead with sterile seawater and held in laboratory conditions for the duration of the challenge. In the main *F. halioticida* challenge used for the analysis (i.e., Challenge 3), there were zero mortalities in the control. Mantle tissue samples were taken during lethal sampling following the death of the individual in the trial or the end of the trial. Individuals that survived to the end of the trial were given a time-to-event value of six to differentiate from the scallops that died on the last day of the trial, and this time-to-event phenotype was used for survival analysis (see below). Tissue samples were stored in 95% ethanol until DNA extraction. Genomic DNA was extracted as described above, following the tissue protocol.

The *F. halioticida* disease trial was analyzed as either binary (alive or dead) and time-to-event (days) within R (R Core Team, 2026) using a Kaplan-Meier survival model to estimate and visualize survival probabilities for each scallop family over time using *survival* (Therneau and Grambsch, 2000; Therneau, 2024) and *survminer* (Kassambara et al., 2024). Families with fewer than five individuals in the disease trial were removed from these analyses, and families that were determined to be potentially contaminated with other families (see *Results* section 3.5) were also removed (i.e., families F4, F9, F18). Significant differences in survival between families were determined by a parametric regression model with a log-logistic distribution to estimate Accelerated Failure Times (AFTs) for each family. Pairwise comparisons of family-level AFTs with Bonferroni multiple test adjustments were performed using *emmeans* (Lenth, 2024). An ANOVA followed by Tukey’s HSD post-hoc tests with Bonferroni multiple test correction was used to test for differences in size between scallop families used for the disease challenge. Model fits were assessed by comparing Akaike Information Criterion (AIC) values between model distributions and by visualizing residual variance and Q-Q plots in base R or with *DHARMa* (Hartig, 2024). Scallop survival and size boxplots were generated with ggplot2 (Wickham, 2016).

Weathervane scallops were purchased commercially from a fishmonger indicating the source was wild-caught weathervane scallops from an Alaskan fishery. Small sections of tissue were stored in 95% ethanol until DNA extraction. Genomic DNA extraction was conducted by Monarch genomic DNA extraction kits.

To compare between non-lethal tissue sampling approaches to use for genomic DNA extraction, DNA was either sampled using the hemolymph extraction method by syringe (see above) or by allowing the scallops to sit in 500 ml beaker overnight, accumulating pseudofaeces, then filtering the water using a 0.22 µm screen and extracting from filters using Monarch genomic DNA extraction kits.

For all samples listed above, DNA was quantified by spectrophotometry using a µCuvette® G1.0 (Eppendorf) measured on a BioSpectrometer (Eppendorf) and normalized to a minimum of 20 ng/µl when concentrations allowed. Samples with too low concentrations were still included in the study at the highest concentration possible. Samples were put in haphazard order by starting concentration, plated in eight 96-well plates, and submitted for genotyping in two separate submissions to the Thermo Fisher Scientific Laboratories in Austin Texas as part of the pilot program for AmpliSeq genotyping. A blank negative control sample was placed in a randomized position on each plate.

### 2.4 Genotyping, filtering, and quality control

Additional sample normalization, library construction and sequencing were conducted using the standard workflow for the AgriSeq amplicon panels on an Ion Torrent S5 instrument (Thermo Fisher). In brief, scallops were genotyped as per the AgriSeq HTS library kit user guide, by using multiplex PCR with the Myes_v.1.0 panel. Each sample had its own barcode for individual identification, and the prepared library was sequenced using two Ion Torrent S5 sequencing chips. Multiple sequencing attempts were made in an attempt to recover low concentration samples, and the run that provided the highest coverage as estimated by total file size was included in the analysis. Results were provided as Torrent VariantCaller tab-delimited text files as well as demultiplexed fastq datafiles.

Torrent VariantCaller output data analysis (i.e., hotspot SNPs) was performed using the *amplitools* pipeline (Sutherland et al., 2024). Demultiplexed fastq files were used for *de novo* SNP calling as per Thompson et al. (2025). The Thermo Fisher AgriSeq amplicon workflow defines target regions as those genomic regions that are amplified and sequenced, and hotspot SNPs as known target SNPs within the target regions. Hotspot (target) SNPs were characterized and converted to a genind file using custom scripts within the *amplitools* pipeline, retaining samples with a genotyping rate (GR) > 70%, retaining SNPs with a GR > 70%, and removing monomorphic loci. SNP calling (*de novo*) was conducted by aligning all raw fastq datafiles from the two sequencing runs against the chromosome-level reference genome using *bwa mem* (Li, 2013) adding read group identifiers unique to each sample (flags: -k 19 -c 500 -O 0,0 -E 2,2 -T 0). SAMtools (Danecek et al., 2021) was used to convert from SAM to BAM format, filtering for quality of alignments (flags: -q 1 -F 4 -F 256 -F 2048), and to sort BAM files. Sorted BAM files were merged into a single file using SAMtools merge. Genotyping was conducted using bcftools mpileup and call (Danecek et al., 2021), setting maximum per-file depth of 230,400 reads, which corresponds to a maximum of 300x coverage in all 768 samples. Samples were then renamed in the raw BCF file using bcftools.

Variant filtering was conducted in a stepwise process using bcftools using the custom script filter_bcf.sh of *amplitools*, evaluating the numbers of variants passing each filter through the steps. The following filters were applied: (1) remove variants within 5 bp from an insertion/deletion (indel); (2) remove variants that are missing in more than 15% of individuals; (3) retain SNP variants only; (4) retain SNPs with a quality score of at least 99 in at least one individual; (5) retain SNPs with an average depth across samples of at least 10 reads; (6) retain biallelic SNPs only; and (7) set any genotype with fewer than 10 reads, more than 10,000 reads, or with a genotype quality score under 20 to missing, then remove variants that are missing in more than 15% of individuals. The output BCF file was converted to a VCF file using bcftools and used for downstream analyses. Target SNPs and *de novo* identified SNPs were plotted along chromosomes using Circos (Krzywinski et al., 2009).

Population genetic analysis used the functions available from *simple_pop_stats* (see Data Availability) using R. The VCF file was read into R using vcfR (Knaus and Grünwald, 2017) and converted to genind format (Jombart and Ahmed, 2011). Populations were assigned to each sample in the genind object and per sample missing data was calculated. Any samples with genotyping rates lower than 70% were removed from the analysis. All code to run the analysis is provided (see Data Availability, *ms_scallop_panel*).

### 2.5 Parentage analysis

The genind object was subset into all separate populations then repooled using *adegenet* to retain only the outplanted (farm) offspring and the 2021 and 2022 broodstock (i.e., SFR2-SFR4 broodstock). Genetic relatedness between individuals was calculated using *related* (Pew et al., 2015), and putative identical individuals were removed (some broodstock individuals lost physical identification tags and had new tags applied, resulting in unexpected identical DNA samples with different identifiers). All broodstock individuals were grouped into a single broodstock population. Monomorphic loci were removed from the dataset. The data were converted from genind format to rubias format (Moran and Anderson, 2019) and exported as a tab-delimited text file specifying offspring and parental generations.

The rubias format text file was read into R and analyzed using the *ckmr_from_rubias* function of *amplitools* to conduct CKMR-sim simulations of expected log-likelihood values for parent-offspring, full-sib, half-sib, and unrelated pairs, and to subsequently generate log-likelihood values for offspring against potential parents (Anderson, 2024). Relationship networks were then constructed using the iGraph toolkit (Csárdi et al., 2024). Both the hotspot and *de novo* datasets were used to simulate expected distributions of the different relationships, but only the *de novo* SNP data were used for the full parentage analysis.

### 2.6 Low-resolution GWAS and heritability for survivorship to *F. halioticida* infection

The full dataset was subset to only include samples from the *F. halioticida* trial, and monomorphic loci and individuals without recorded survival phenotypes were removed. A principal components analysis (PCA) was conducted using *adegenet* to confirm family groupings for quality control of sample labels. The original VCF file was then subset to only include the retained samples and converted to the required input files for GEMMA (Zhou and Stephens, 2012) using custom code (see Data Availability, *ms_scallop_panel*). A GWAS using survival as a binary phenotype was conducted in GEMMA using loci with MAF > 0.05, and a Manhattan plot was generated using fastman (Paria et al., 2022).

To evaluate heritability of the survival phenotype, the VCF file with only the samples from the *F. halioticida* trial was read into R using vcfR (Knaus and Grünwald, 2017) alongside phenotypes. The genotypes were converted from two-character allele format to dosage of the alternate allele (0,1,2), and the *qc.filtering* function of ASRgenomics (Gezan et al., 2022) was used to obtain an M-matrix, which was then converted to a G-matrix. The function *kinship.diagnostics* was used to identify putative duplicate individuals or extreme values, and the G-matrix was converted to an inverse G-matrix using the function *G.tuneup* with the bend argument enabled. A genomic best linear unbiased predictor (gBLUP) was run using MCMCglmm (Hadfield, 2010) using a ‘threshold’ trait distribution with binary survival as the response variable and individual size as a fixed effect, and the individual as a random effect, with 400,000 iterations (30,000 burn-in, and 5 iteration thinning). The heritability was calculated as the 95% confidence interval and posterior mean of the individual variance component divided by all variances.

### 2.7 Genetic differences among weathervane scallop and Yesso scallop from Japan and Canada

To evaluate the ability of the panel to amplify and differentiate Yesso scallop from weathervane scallop, data were obtained from a variety of sources. Reduced-representation sequencing (i.e., ddRAD-seq) fastq files were obtained from Sutherland et al. (2023) and 20 samples were selected haphazardly from each of Japan wild, BC farm, and the VIU breeding program. Amplicon panel fastq files were obtained from the present pilot study and 10 samples were selected haphazardly from the 2022 broodstock and 10 samples from the 2022 spawn event offspring. Weathervane samples were genotyped only by the amplicon panel, and all available samples were used (n = 6).

To facilitate comparability between platforms, a common genotyping approach was taken. The ddRAD-seq and amplicon panel fastq files were used together as an input to the *de novo* genotyping workflow of *amplitools*, using the same approach and filtering parameters as described above, except requiring a 90% GR per locus to include the SNP in the analysis. Filtered genotypes were analyzed in R using *simple_pop_stats* as described above, removing any individuals with genotyping rates under 70%. The genotyped samples were analyzed in an unsupervised PCA and a supervised discriminant analysis of principal components (DAPC) to best separate the collections based on population labels. The number of fixed differences in genotypes between all pairs of collections in the analysis were inspected using the *gl.fixed.diff* function of dartR (Gruber et al., 2018). Samples were clustered and visualized using the heatmap function in R based on allelic dosage.

## 3 Results

### 3.1 Panel design, sequencing, and *de novo* genotyping

There were 9,331 SNPs from a Yesso scallop ddRADseq study (Sutherland et al., 2023) with positions on the newer, chromosome-level assembly (Liu et al., 2020). Of these, 3,057 passed additional filters (i.e., MAF > 0.01; no significant deviation from Hardy-Weinberg proportions in collections from Japan or VIU). Limiting the dataset to only those SNPs with *H*_OBS_ < 0.5 in each of the two collections resulted in 2,821 retained SNPs. Marker selection picked SNPs from 50 non-overlapping 20.1 Mbp windows to obtain the most SNPs with highest heterozygosity (Table S1; Table S2). The 592 SNPs selected by this approach were increased to 610 by adding additional SNPs with high *H*_OBS_ regardless of genomic position. The selected SNPs in the source ddRADseq samples had an average *H*_OBS_ of 0.37 and 0.36 in the Japan and VIU collections, respectively.

Of the 610 selected SNPs, 601 had 200 bp flanking both sides of the SNP that could be extracted for primer design. Four SNPs were removed at this stage due to the presence of more than two alleles at the target SNP. Two SNPs associated with shell colour on chr11 were added to the design (SNP_6363 and SNP_259; Zhao et al., 2017), resulting in 599 target SNPs (Additional File S1). Following the panel design, each chromosome of the assembly had an average (± s.d.) of 29.5 ± 17.1 SNPs (range: 16-99 SNPs; Figure 1). The design was successful for 587 target SNPs present in 579 amplicons.

**Figure 1.**
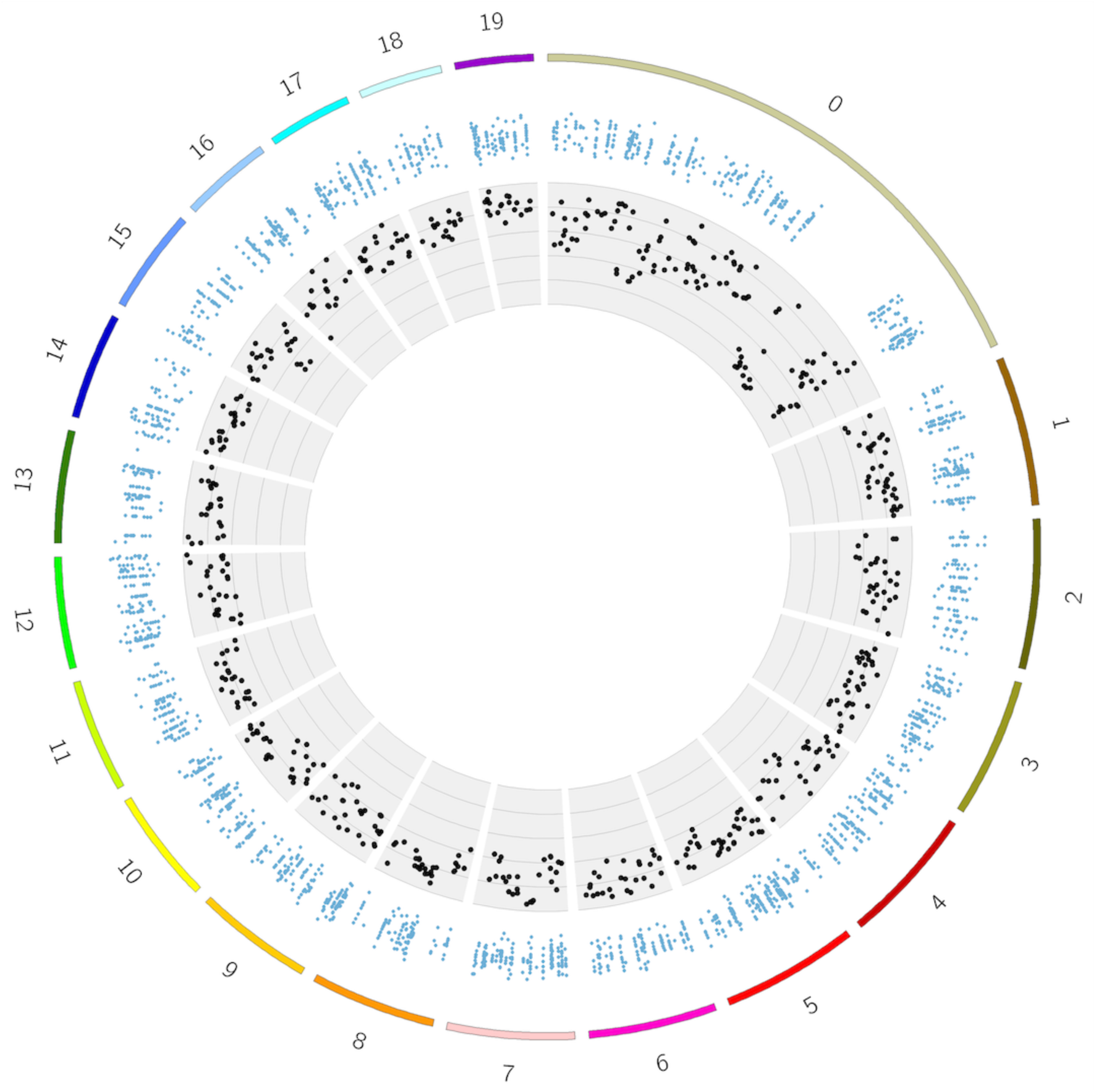
Yesso scallop chromosome-level genome assembly plotted with 19 chromosomes shown with hotspot target SNPs (black points) shown as cumulative observed heterozygosity (summed for Japan and Canada ddRADseq source data). Also shown are the positions of the *de novo* identified SNPs (small blue points, with a vertical jitter for ease of display). Note: the reference genome used has all remaining scaffolds joined in Chr00, which is also shown here.

**Figure 2.**
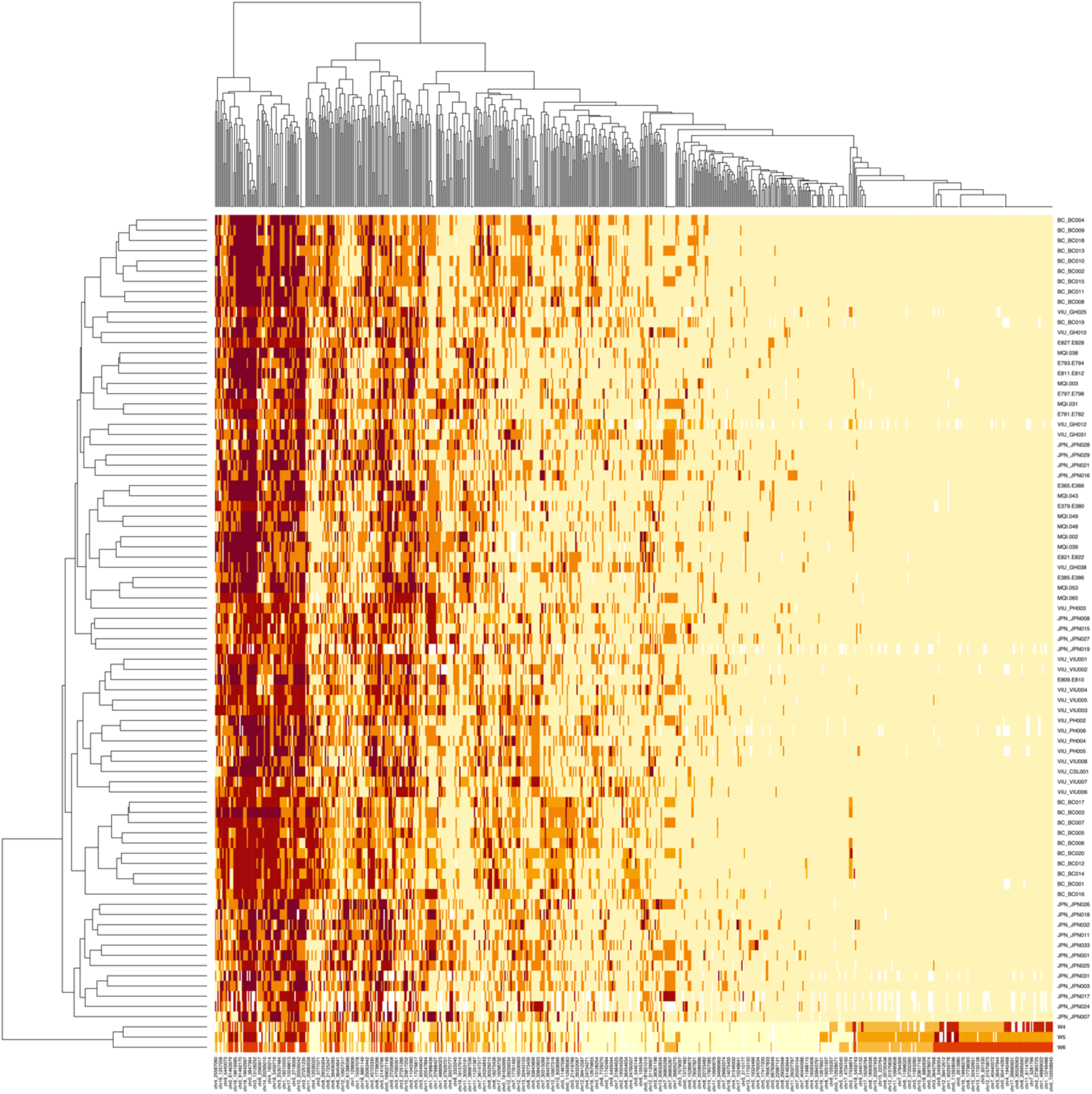
Genotypes of individuals (rows) plotted as a heatmap (yellow = 0/0; orange = 0/1; red = 1/1) for all loci (columns) in the Yesso scallop vs. weathervane scallop analysis. Clustering was enabled for both rows and columns, showing that the weathervane samples (n = 3) clustered separately at the bottom of the heatmap, with a clear set of loci that are only polymorphic in the weathervane scallops (bottom right). A high degree of shared polymorphism is present in the two species.

### 3.2 Genotyping *de novo* and target (hotspot) SNPs

In total, 171,956,927 reads were obtained from the 759 individuals in the pilot study, with an average of 226,557 ± 275,073 reads per sample (median: 142,273; range: 11-2,530,237; Additional File S2). Alignment rates against the genome were on average 94.1 ± 3.0% per sample (median: 95.0%). Genotyping the amplicons to identify SNPs *de novo* identified 2,983,897 raw putative variants, and after filters were applied (see *Methods*), 2,592 SNPs were retained from throughout the 19 chromosomes and additional scaffold (Figure 1; see Table S3). The blank samples (n = 9) had an average of 6.4k ± 6.3k reads each (range: 1.5k-22.1k reads; Additional File S2).

The main pilot study experiments that were used to evaluate panel performance (see Table 1), consisted of a parentage experiment (post-filters: n = 76 potential parents; 61 offspring) and a low-resolution family-based GWAS and heritability evaluation for survivorship to *F. halioticida* exposure (n = 431 offspring). The *de novo* analysis resulted in the retention of 716 samples after filtering (Table 1). The broodstock from the 2022 spawn event had higher genotyping rates (GR) than those from the 2021 spawn, with 95.4% and 86.9% of individuals retained following GR filters, respectively (Figure S1A). All outplanted offspring passed GR filters; these samples were taken using a syringe to obtain a small volume of hemolymph. The tests of genotyping filters following flow-through of scallop holding water and pseudofaeces were not successful, as the samples had an average GR of 27.4% (Table 1; Figure S1A). Amplification of non-target weathervane scallops (*P. caurinus*) resulted in half (n = 3) of the weathervane samples passing GR filters, and low sequence coverage overall for weathervane samples was observed (36,113 ± 33,373 reads per sample; median = 31,819 reads).

Analysis with the target (hotspot) variants (see *Methods*) resulted in 581 retained SNPs and 766 individuals before filters were applied, with an average missing data per sample of 16.1 ± 18.5%. Retaining samples with GR > 70% resulted in the retention of 683 individuals. When the hotspot data were subset to only include the *F. halioticida* trial, and broodstock and offspring from the VIU breeding program, an additional 34 monomorphic SNPs were removed, and 44 SNPs were removed due to low GR (n = 503 SNPs retained). In almost all the collections, the number of retained samples was greater in the *de novo* dataset than the hotspot dataset (Table 1), suggesting that the *de novo* approach not only resulted in more loci being genotyped but also more individuals being retained for analysis.

### 3.3 Parentage study

The panel was evaluated for use in parentage studies by identifying the parents of outplanted offspring from the VIU breeding program that were produced in the 2022 spawn events (SFR3 and SFR4). All broodstock used in 2021 and 2022 spawn events were included in the analysis. This analysis included 61 offspring and 76 potential broodstock (using *de novo* SNPs). Two putative replicate broodstock DNA samples were identified and removed (see *Methods*). After monomorphic loci were removed from the parentage dataset, 1,863 SNPs were retained. Using a parallel approach, the target (hotspot) SNP dataset retained 503 SNPs. Simulations with CKMR-sim indicated a strong ability to differentiate parent-offspring or full-sib relationships from unrelated pairs using either the hotspot data (Figure 3A) or the *de novo* SNPs (Figure 3B), but the *de novo* SNP dataset was estimated to have increased resolution relative to the target (hotspot) dataset.

**Figure 3.**
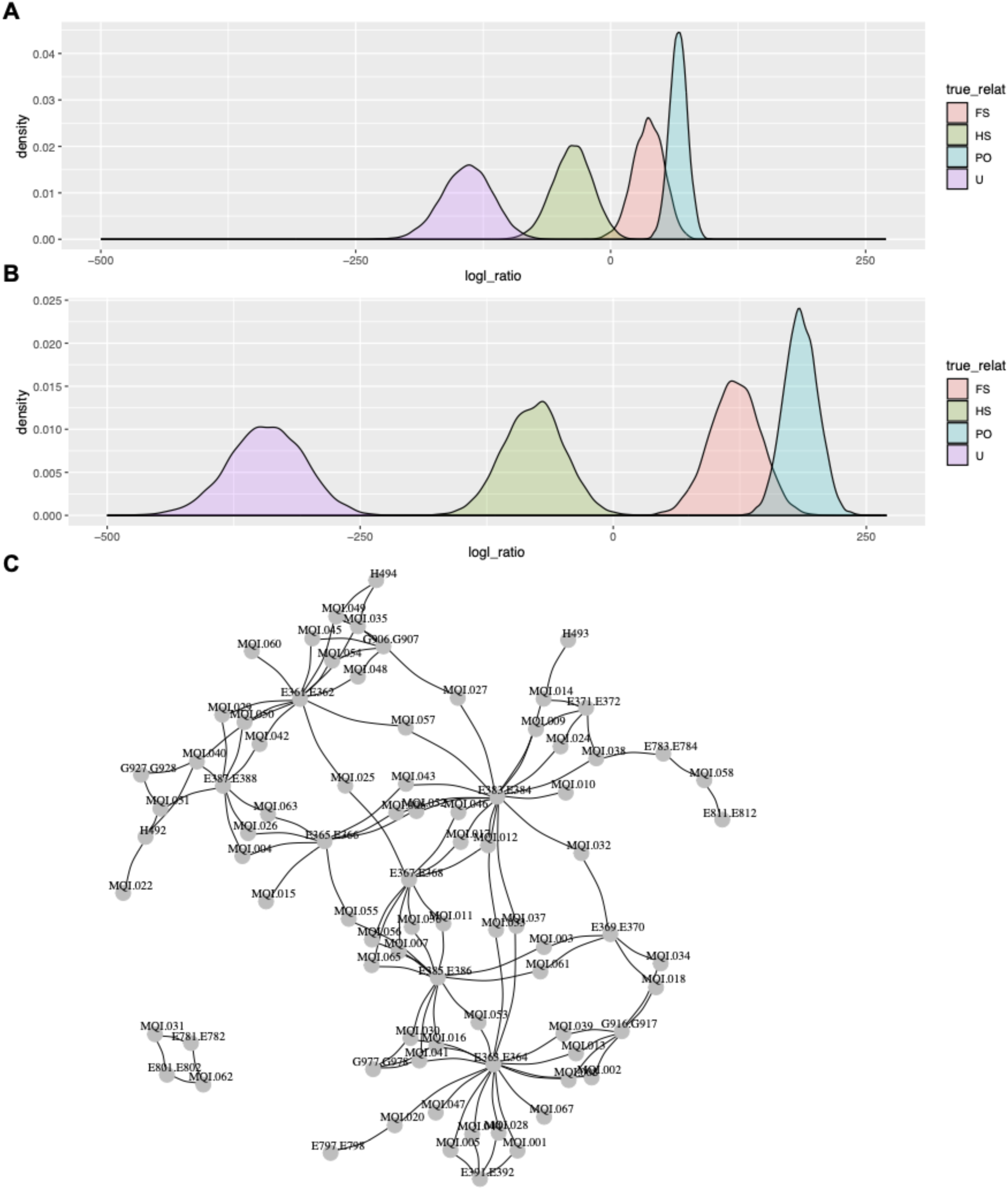
Evaluation of panel for parentage showing CKMR-sim log likelihood distributions for simulated parent-offspring (PO), full-sib (FS), half-sib (HS), and unrelated (U) pairs of individuals using genotypes based on (A) the target design (hotspot) SNPs (n = 503 SNPs); and (B) *de novo* identified loci (n = 1,863 SNPs). (C) Identified empirical relationships in the data in an iGraph network plot showing links between putative parents and offspring (log likelihood ≥ 20; offspring denoted with ‘MQI’ identifier).

Parentage assignments were conducted then compared with documented spawn designs for SFR3 and SFR4. Two-parent assignments were made using parent-offspring relationships for 47 of the 61 offspring (77.0%), and an additional 11 offspring were assigned to family based on sibship assignments to sibs with identified parents, resulting in 58 of 61 offspring (95.1%) assigned to family (Table 3; Figure 3C; Additional File S3). Five of the assigned offspring were assigned to the same parental pair that was used in both SFR3 and SFR4, and so the spawn year was not determinable for these offspring (note: Yesso scallop are iteroparous and spawning was non-lethal). The remaining three offspring only received a single-parent assignment or assignments to parents of the same sex (i.e., false positive secondary assignments), and therefore were not unambiguously assigned to a family.

**Table 3.**
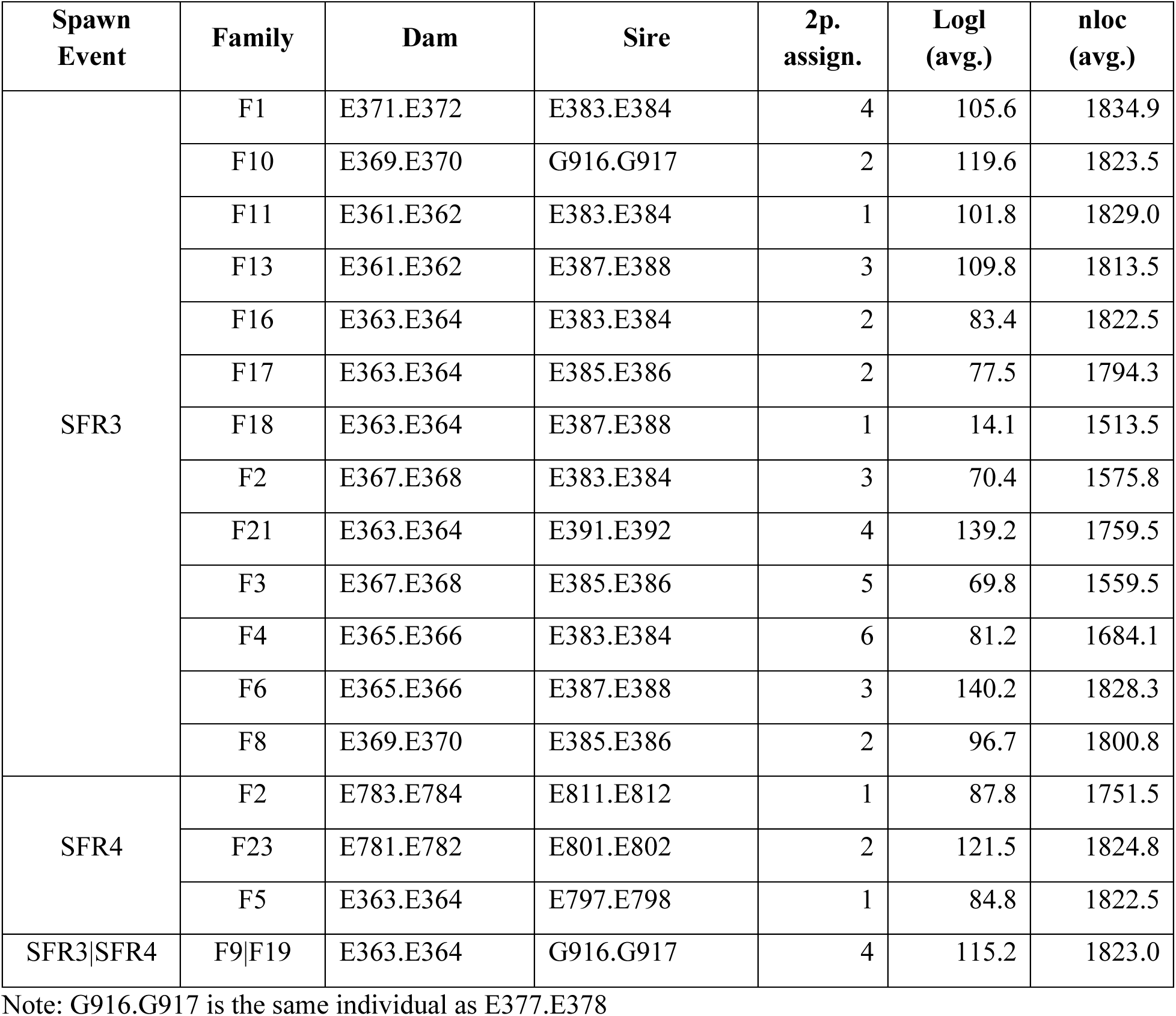
Parentage assignment summary for VIU outplanted samples including the spawn event, family, dam and sire, as well as the number of offspring that received two-parent assignments (2p. assign.) and the average log likelihood and number of loci used in each assignment. Offspring that were not assigned or that required sibship analysis to determine family were not included here but are shown in Additional File S3. One combination of dam and sire was used in both spawn events and therefore is unknown which spawn event from which it originated (i.e., F9/F19). SFR = scallop family run; F = family.

### 3.4 *F. halioticida* challenge with low-resolution GWAS and heritability estimate

Survivorship to a *F. halioticida* challenge was analyzed over a five-day period in 10 full-sib families, and significant differences in survival probabilities were observed between families (p < 0.01; Figure S5). Specifically, higher survival probabilities were observed for families F8, F12, and F17 compared with families F1 and F5 (Figure S5; Figure S6; Additional File S4). Notably, there were also significant size differences between families (Additional File S4), where family F1 was the smallest, whereas F16 and F12 were the largest (Figure S6B; model assumption plots in Figure S7). Size correlated with survival; the families with the largest scallops typically had higher survival (Figure S6C). However, this was not always the case, for example, although F8 and F17 had significantly higher survival than F5, they were not significantly different in size (Figure S6B; Additional File S4). Notably, there was substantial intra-family variance in both phenotypes.

Selective breeding for *F. halioticida* resistance could be possible using marker-assisted selection, if significant large-effect loci are identified, or it could be possible using genomic selection (GS), if the trait is sufficiently heritable. The *F. halioticida* exposure trial dataset contained 405 individuals with survival phenotypes, and these samples contained 1,599 polymorphic SNPs (*de novo* genotyping). On average, 22.5 individuals were genotyped from each family (s.d. = 31.3; range = 1-141; Table S4). Families generally grouped in a PCA clustering samples by genotypes (Figure 4A), with several notable exceptions. A subset of samples labeled as families F4 (n = 10), F9 (n = 23; from both challenge 3 and 4), and F18 (n = 2) were grouped within the F8 cluster in the PCA, suggesting that these three families were contaminated at some point in the hatchery or exposure trial with individuals from family F8 (i.e., genetically the samples are from family F8, contrary to their metadata labels). Family F9 was challenged twice (i.e., challenge C3 and C4), and parentage confirmed the parents for the two trials (aside from the F8-contaminated samples described above). Consistent parents were also confirmed for family F19 samples held in two separate tanks (i.e., B2 and B8), allaying hatchery concerns of mixed up identities of one of the tanks. Given that the downstream GWAS and GS analyses do not depend on family identifiers as they calculate relatedness based on the genotypes, these findings were noted but changes to metadata was not required for the downstream analyses.

**Figure 4.**
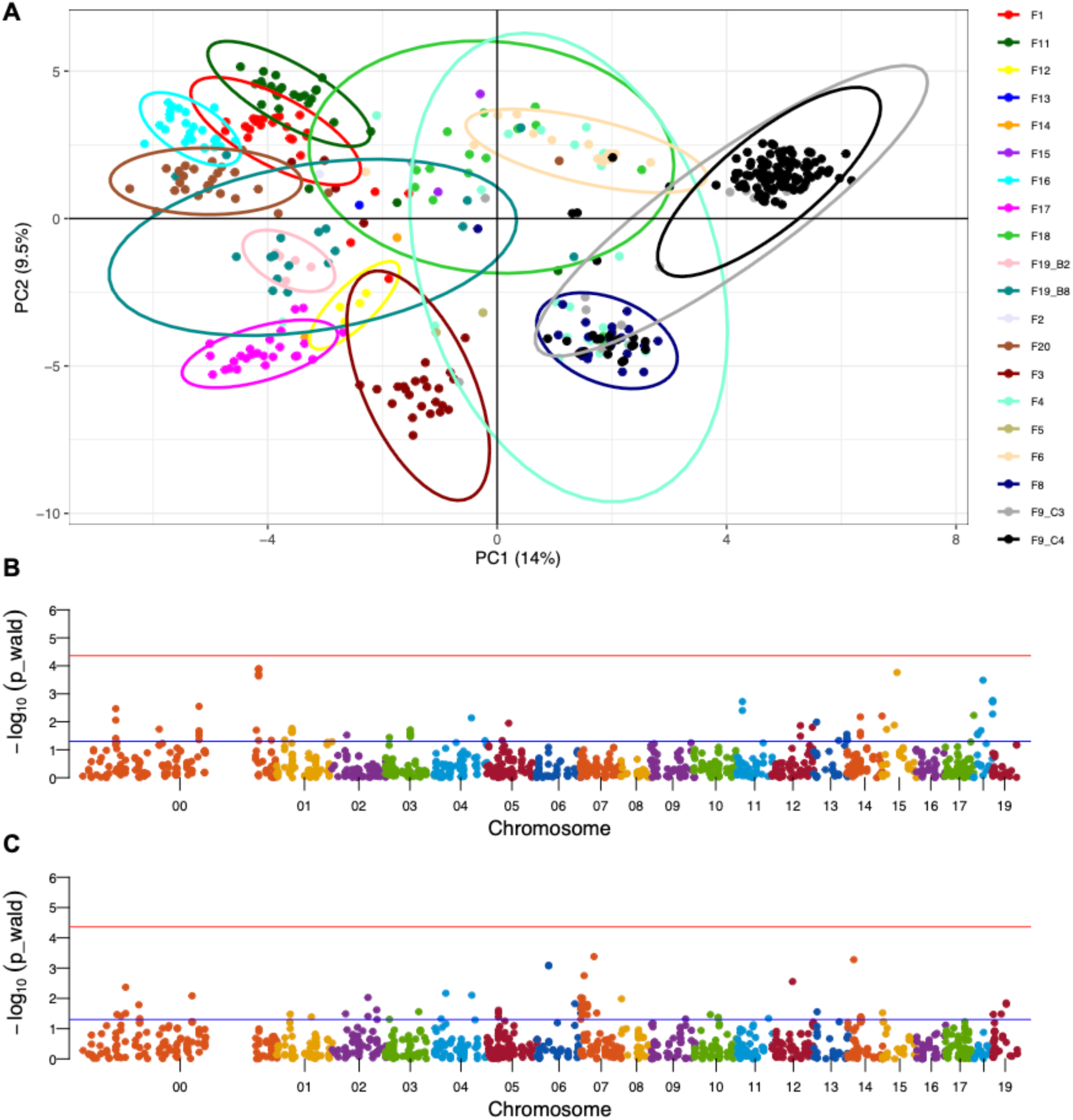
(A) Principal components analysis (PCA) showing samples clustered by genotypes. Family clustering is observed for the 18 unique families included in the *Francisella halioticida* challenge trials. Using *de novo* genotype data, 1,151 analyzed SNPs were inspected for associations to phenotypes (B) survivorship (binary) in the challenge trial; and (C) size of scallop individual at the start of the trial. No significant associations were identified following Bonferroni correction (alpha = 0.05; red line). The blue line shows uncorrected *p* = 0.05.

To investigate whether any genomic segments were associated with survivorship, a GWAS was conducted analyzing all 405 samples from the 10 families using 1,151 SNPs (MAF > 0.05). Survival as a binary trait nor size at the start of the trial showed significant associations at the inspected loci (Figure 4B and Figure 4C). Due to the lack of any large effect regions or loci being associated with survivorship, the heritability of the trait was estimated to determine whether a genomic selection approach could be used for the trait. The model of genotypic variance showed a correlation between size and binary survival, and therefore the applied model used size as a fixed effect. The model passed the stationarity test, and size and intercept were both found to be significant fixed effects (p < 0.0001). The narrow-sense heritability for the model was calculated as 0.11 (95% CI = 0.002-0.302), suggesting a low but non-zero heritability for the trait.

### 3.5 Weathervane scallop genotypes and comparison to Yesso scallop in BC and Japan

Several weathervane scallops were amplified using the Yesso scallop panel to test cross-species amplification and species differentiation with the selected markers. It was also used to determine whether there was any evidence for a past hybridization event between the BC scallops analyzed and the weathervane scallop (see *Methods* for sample sources; Additional File S2). The Yesso scallop ddRADseq data used for the analysis had an average (± s.d.) of 2.47 ± 1.22 M reads per-sample (alignment rate = 79.1 ± 9.1%) and the Yesso scallop amplicon panel data had an average of 0.34 ± 0.26 M reads per-sample (alignment rate = 95.4 ± 1.1%). The weathervane scallop amplicon panel data had an average of 0.04 ± 0.03 M reads per-sample (alignment rate = 88.0 ± 2.4%). After all samples were aligned to a common reference genome, genotyping was conducted with 125,766,345 read alignments. This identified 445,934 raw variants, 686 of which passed all filters (Table S3). Filtering individual samples based on GR retained all samples except three weathervane samples and one VIU ddRADseq sample (n = 82 samples retained; Figure S1B).

The greatest source of the variation in this dataset was between the Yesso scallops and weathervane scallops, as shown in a PCA separating the two species on PC1 (PVE = 9.6%; Figure S2A). PC2 (PVE = 6.9%) separated samples based on sample source and technology, with the VIU samples genotyped by ddRADseq and those by amplicon panel grouping separately. PC3 captured the variation separating Japan Yesso scallops from the rest of the samples (PVE = 5.6%; Figure S2B). A DAPC showed similar trends, with discriminant function 1 (DF1) separating the weathervane scallops from the Yesso scallops, and a smaller extent of difference between genotyping platforms (Figure S3).

Polymorphism density was estimated for each population. The weathervane scallops (n = 3), were isolated and monomorphic loci were removed. This left a total of 52.0% of the retained loci being polymorphic (n = 357 loci). For comparison, Yesso scallops from BC or Japan were randomly subsampled to three individuals and monomorphs were removed, resulting in an average (± s.d.) of 45.0 ± 2.3% and 45.3 ± 1.8% polymorphic loci, respectively (Figure S4).

The extent of genetic differentiation between each Yesso scallop collection and the weathervane scallop collection was first investigated by identifying fixed genetic differences (i.e., weathervane vs. Japan wild (ddRADseq) or VIU (ddRADseq or panel)). No fixed differences were observed between any of the Yesso scallop collections, and an average of 17.5 loci had fixed differences between the Yesso collections and the weathervane scallops (Table 2). Both the wild Japan and VIU breeding population Yesso scallops sequenced by ddRADseq had 17 loci with fixed differences observed when compared with the weathervane scallops. To visualize the extent of shared or species-specific polymorphism, a heatmap was generated based on genotypes (i.e., 0, 1, or 2 alternate alleles) per locus, per individual (Figure 2). Although many loci show shared polymorphism between the species, there are clear sections of the heatmap with polymorphism specific to weathervane or Yesso scallops. Based on these analyses, the VIU breeding population did not appear to be genetically more similar to the weathervane scallop than were the Japan wild population, and therefore, no evidence of a past hybridization event was observed in these samples.

## 4 Discussion

The present study demonstrates the utility and versatility of the newly-designed Yesso scallop low-density genotyping amplicon panel. The design methodology targeted markers with high heterozygosity in both Japan and Canada, but given the high SNP density of Yesso scallop as species, the panel is likely to be useful in any population using the *de novo* genotyping approach, as long as the primers anneal and amplify in the non-target population. This target site flexibility is one of the benefits of amplifying genomic regions instead of genotyping specific SNPs, as discussed for Pacific oyster (Thompson et al., 2025) and of particular relevance to organisms with abundant genetic polymorphism with high turnover rates. By using the *de novo* dataset, an additional benefit was observed as more samples were retained through filters compared to the hotspot target dataset. The panel has been integrated into the *amplitools* framework, enabling the same tools and pipelines as developed for Pacific oyster (Sutherland et al., 2024), including microhaplotype genotyping (Thompson et al., 2025), and imputation for use in genomic selection (Sutherland et al., 2026).

Genotyping of marine bivalves to accelerate breeding program progress is made possible by high-throughput SNP-based methods. The choice of platform is driven mainly by cost, required marker density, and need for cross-study comparability. Common genotyping platforms include SNP arrays (Gutierrez et al., 2017; Guo et al., 2023), reduced-representation sequencing (e.g., RAD-seq, DArTseq; Vu et al., 2021), low coverage whole genome resequencing and imputation (Yang et al., 2024a), and amplicon panels (Meek and Larson, 2019; Sutherland et al., 2024). SNP arrays are highly reproducible and have moderate costs (approx. $30 to $70 USD per sample depending on SNP density), but the high genetic polymorphism and turnover rate, as well as structural variation in marine bivalves (e.g., large deletions) can cause probe failure and significantly reduce call rates (Guo et al., 2023; Ajithkumar et al., 2025). Reduced-representation sequencing uses restriction enzymes and size selection to limit sequencing to regions around specific cut sites, providing a cost-effective approach for some projects (e.g., $30 USD per sample), but may lack locus consistency across studies and laboratories (Vu et al., 2021). Low-coverage whole genome resequencing involves sequencing the entire genome at a very shallow depth, which typically precludes accurately calling of individual genotypes, but genotype likelihoods can be used for population-level metrics or comparisons (Lou et al., 2021). It is also possible to combine low-depth resequencing with imputation to estimate individual-level genotypes (Yang et al., 2024a). Whole-genome methods provides extensive genomic coverage and allows the identification of millions of SNP variants, but also may come with additional bioinformatics requirements. Moderate depth genome resequencing (e.g., 10X coverage) can provide reliable individual level genotypes but is often more expensive, with costs scaling with genome size (e.g., $50-100 USD per sample) (Sutherland et al., 2026), but also see Yang et al. (2026) for cost reduction methods for library preparation. Given the above, the developed amplicon panel was built with the aim to provide an inexpensive (approx. $14 USD) and consistent method of reliably genotyping the same set of loci across samples with low bioinformatic requirements (Meek and Larson, 2019) and increased flexibility for target SNPs and ability to genotype with lower quality or lower quantity input DNA (Sutherland et al., 2026) that can be used for parentage and genomic selection (Delomas et al., 2023; Sutherland et al., 2024; Thompson et al., 2025).

The parentage and sibship analyses conducted here demonstrate the panel’s utility for supporting breeding program progress. For example, a specific family (F8) was found to have contaminated three other families (now mixed families); these three families should be excluded from directional selection unless potential broodstock are genotyped and confirmed to be from the target pedigree line. This is an empirical documentation of the risk of family contamination, which carries potential for inhibiting breeding gains, as discussed by Hedgecock and Davis (2007). This finding led to the removal of these mixed families in the analysis of family-based survival differences in response to *F. halioticida*, and therefore reducing the noise in the analyzed dataset. The GWAS and genomic selection heritability estimate analyses were able to use these samples as family identifiers are not used, but rather relatedness is calculated directly from the genotypes.

*Francisella halioticida* has emerged as a significant global pathogen of farmed marine bivalve in North America (Meyer et al., 2017), Asia (Kawahara et al., 2018) and Europe (Charles et al., 2020; Cano et al., 2022). The present work is the first to report significant family-level differences in resistance to *F. halioticida*. This finding is highly valuable to the aquaculture industry, as farmers are actively trying to limit their losses to this disease. The experimental infection conducted here suggests that the mortality observed was due to *F. halioticida*, given the absence of mortality in the controls. However, a limitation of the study should be noted in that there were inter-family differences in scallop sizes, and size was found to correlate with survival in the trial (see Results). It is possible that the reduced survivorship in some families with smaller individuals may be related to dosage of *F. halioticida*, as all individuals received the same bacterial injection volume. Future exposures could normalize input volumes based on size, although this will increase the labour required. Importantly, there were also families that had significantly higher survival without significant differences in size, suggesting that there is likely a combination of factors, including genetics, that resulted in increased survival.

Genetic tools are needed to increase disease resilience in farmed shellfish. Shellfish farming industries are unique with regards to disease management as there are very few ways to reduce the detrimental impact of pathogens on production (Pernet et al., 2016). Marine bivalves are typically farmed in estuaries and coastal bays, which strongly precludes the use of chemical treatments due to the quantity of product required and the potential impacts on the local environment. Consequently, the industry has relied upon pedigree-based genetic selection programs to produce disease resistant stock, and this has been successful for a number of oyster pathogens (reviewed by Dégremont et al., 2015). A growing body of evidence indicates that genomics, including genomic selection, will further accelerate genetic gains for disease resistance in marine bivalves by allowing selection of superior broodstock compared to traditional, pedigree-based methods (Gutierrez et al., 2020; Divilov et al., 2023; Yang et al., 2024b; Ajithkumar et al., 2025; Wang et al., 2025; Jourdan et al., 2026). The present study also provides the first report of SNP-based heritability estimate for Yesso scallop resistance to *F. halioticida*. The estimate of low heritability (*h^2^* = 0.11) indicates that genomic selection for enhanced disease resistance to *F. halioticida* is possible within the VIU breeding program. No major effect loci were observed in the family-based GWAS using *de novo* SNPs from the amplicon panel, and so there is no evidence yet for the potential for marker-based selection. Future GWAS experiments with less related individuals would benefit from using a high-density genotyping platform (e.g., >50,000 SNPs). The present GWAS used families, and so large linkage blocks are expected, as occurs in marine bivalves in family-based breeding programs (Hu et al., 2022); an increase in marker density for the present analysis may not provide additional benefit. Future studies may also benefit from expanding the number of evaluated individuals (e.g., >1,800; Goddard and Hayes, 2009) and the diversity of genetic backgrounds. These changes would increase statistical power to detect moderate to large effect loci, if they exist. Comparisons of low- and high-density genotyping have found that several thousand SNPs are adequate for genomic selection in marine bivalves for growth and disease traits (Gutierrez et al., 2018; Vu et al., 2021; Delomas et al., 2023). Polygenic architecture for *F. halioticida* resistance would not be surprising given the many other marine bivalve studies that report polygenic architecture for disease resistance to bacterial or parasitic diseases (Yang et al., 2024b; Ajithkumar et al., 2025; Wang et al., 2025).

The cross-species amplification provided the opportunity to compare weathervane scallop and Yesso scallop at the loci genotyped in both species. Using the aquaculture Yesso scallop samples genotyped here, we did not find any evidence of a past weathervane x Yesso scallop hybridization event. Specifically, there was no increase in genetic similarity observed between BC (cultured) or Japanese (wild) Yesso scallops compared to Alaskan weathervane scallops. If a signal of hybridization was detected, we would have expected fewer fixed differences in the weathervane-BC aquaculture comparison than the weathervane-Japan wild comparison, and closer clustering of weathervane to BC aquaculture samples than weathervane to Japan wild. Demographic inference and analysis of past hybridization events could be further explored through whole-genome approaches, but we have yet to observe any evidence for hybridization in the cultured BC Yesso scallops examined. It could also be of benefit to analyze additional weathervane scallops from a region more geographically proximal to the likely source of documented past hybridization events, although low genetic differentiation and a lack of isolation by distance is expected over large geographic distances (Gaffney et al., 2010). Notably, there is extensive shared variation that is observed between all analyzed populations of Yesso scallops and weathervane scallops. The amplicon panel may be used for species identification purposes to differentiate between weathervane and Yesso scallop, as has been applied in other species (Beacham and Wallace, 2020), but the reliability of this assay would be improved by genotyping additional weathervane samples.

## 5 Conclusion

This study demonstrates the utility and versatility of a low-cost, genotyping amplicon panel for Yesso scallop breeding applications. Through the use of the panel, family errors in a pedigree-based breeding program were identified and removed to reduce analysis noise and directional breeding inhibition. It was also possible to evaluate whether the existing breeding program showed any large effect genomic segments associated with resistance to *F. halioticida*, and although there were no associations observed, the potential for genomic selection was demonstrated by identifying a low (non-zero) heritability estimate for the trait. An analysis of weathervane scallop also genotyped with the panel did not find evidence of a past Yesso scallop *x* weathervane scallop hybridization within the tested samples. Collectively, this work and provided genetic tool will help the aquaculture industry make best use of the standing genetic variation in Yesso scallop broodstock lines, support genetic research into trait architecture, improve selective breeding progress, and guide hatchery decisions.

## Supporting information

Supplemental Results

## Data Availability

Marker data were obtained from Sutherland et al. (2023): doi.org/10.6084/m9.figshare.22670626

Per locus heterozygosity metrics and Hardy-Weinberg statistics were obtained from Additional File S7 of Sutherland et al. (2023).

Raw RADseq fastq data was obtained from BioProject PRJNA947158.

Raw amplicon panel fastq data has been uploaded to NCBI SRA under the accession PRJNA1281189.

The amplicon panel is available through ThermoFisher Scientific as item WGAG23043_VIU, and the hotspot file, regions file, and target genome have been added to the *amplitargets* repository.

The following GitHub repositories supported this work:

Manuscript analyses: https://github.com/bensutherland/ms_scallop_panel

*amplitools*: https://github.com/bensutherland/amplitools

*amplitargets*: https://github.com/bensutherland/amplitargets

## Additional Files

**Additional File S1.** Marker information provided to the panel design commercial provider, including the marker identifier, source contig-level chromosome identifier and position, and the reference and alternate alleles, as well as the design window (401 bp centered on the target SNP).

**Additional File S2.** Read and alignment details for all samples included in the pilot study for *de novo* SNP discovery and genotyping, and in the analysis of Yesso scallop and weathervane scallop by amplicon panel and RAD-seq genotyping.

**Additional File S3.** Per-individual parentage assignments including the dam, sire, family, and whether the assignment was conducted by parent-offspring relationship or if it required sibship analysis.

**Additional File S4.** Pairwise statistical test results for survivorship or size differences between families.

## Acknowledgements

Thanks to Claudio Carrasco and Dr. Srinivas Chadaram for their work at Thermo Fisher Scientific in supporting the design of the amplicon panel and running pilot samples through the AmpliSeq program. Thanks to Dr. Gary Meyer for providing the *F. halioticida* isolate that was used in the challenge study. Thanks to James Dennis-Orr for support with sampling and tissue collection and to Lauren Krzus for support in scallop spawn events. Thanks to Prof. Christopher Langdon for providing weathervane scallop samples for this study. Thanks to Carl Butterworth of VIU for technical support, and to Provan Crump for conditioning of scallop broodstock. Thanks to five anonymous reviewers and the Editor for comments on an earlier version of the manuscript.

## References

Ajithkumar, M., D’Ambrosio, J., Travers, M.-A., Morvezen, R., Dégremont, L., 2025. Genomic selection for resistance to one pathogenic strain of *Vibrio splendidus* in blue mussel *Mytilus edulis*. Frontiers in Genetics Volume 15 - 2024. 10.3389/fgene.2024.1487807

Anderson, E.C., 2024. CKMRsim: Inference of pairwise relationships using likelihood ratios..

Baetscher, D.S., Clemento, A.J., Ng, T.C., Anderson, E.C., Garza, J.C., 2018. Microhaplotypes provide increased power from short-read DNA sequences for relationship inference. Mol. Ecol. Resour. 18(2), 296–305. 10.1111/1755-0998.12737

Beacham, T.D., Wallace, C.G., 2020. Salmon species identification via direct DNA sequencing of single amplicons. Conservation Genetics Resources 12(2), 285–291. 10.1007/s12686-019-01102-1

Bourne, N.F., 2000. The potential for scallop culture–the next millenium. Aquacult. Int. 8, 113–122.

Bower, S.M., Blackbourn, J., Meyer, G.R., Welch, D.W., 1999. Effect of *Perkinsus qugwadi* on various species and strains of scallops. Dis. Aquat. Org. 36(2), 143–151.

Brevik, O.J., Ottem, K.F., Kamaishi, T., Watanabe, K., Nylund, A., 2011. *Francisella halioticida* sp. nov., a pathogen of farmed giant abalone (*Haliotis gigantea*) in Japan. J. Appl. Microbiol. 111(5), 1044–1056. 10.1111/j.1365-2672.2011.05133.x

Cano, I., Parker, A., Ward, G.M., Green, M., Ross, S., Bignell, J., Daumich, C., Kerr, R., Feist, S.W., Batista, F.M., 2022. First detection of *Francisella halioticida* infecting a wild population of blue mussels *Mytilus edulis* in the United Kingdom. Pathogens 11(3), 329.

Charles, M., Villalba, A., Meyer, G., Trancart, S., Lagy, C., Bernard, I., Houssin, M., 2020. First detection of *Francisella halioticida* in mussels Mytilus spp. experiencing mortalities in France. Dis Aquat Organ 140, 203–208. 10.3354/dao03505

Csárdi, G., Nepusz, T., Traag, V., Horvát, S., Zanini, F., Noom, D., Müller, K., 2024. igraph: network analysis and visualization in R.

Danecek, P., Bonfield, J.K., Liddle, J., Marshall, J., Ohan, V., Pollard, M.O., Whitwham, A., Keane, T., McCarthy, S.A., Davies, R.M., Li, H., 2021. Twelve years of SAMtools and BCFtools. GigaScience 10(2). 10.1093/gigascience/giab008

Dégremont, L., Garcia, C., Allen, S.K., 2015. Genetic improvement for disease resistance in oysters: a review. J. Invertebr. Pathol. 131, 226–241. 10.1016/j.jip.2015.05.010

Delomas, T.A., Hollenbeck, C.M., Matt, J.L., Thompson, N.F., 2023. Evaluating cost-effective genotyping strategies for genomic selection in oysters. Aquaculture 562, 738844. 10.1016/j.aquaculture.2022.738844

DFO, 2023. Aquaculture production quantities and value. https://www.dfo-mpo.gc.ca/stats/aqua/aqua-prod-eng.htm.

Divilov, K., Merz, N., Schoolfield, B., Green, T.J., Langdon, C., 2023. Marker-assisted selection in a Pacific oyster population for an antiviral QTL conferring increased survival to OsHV-1 mortality events in Tomales Bay. Aquaculture 567, 739291. 10.1016/j.aquaculture.2023.739291

Dou, J., Li, X., Fu, Q., Jiao, W., Li, Y., Li, T., Wang, Y., Hu, X., Wang, S., Bao, Z., 2016. Evaluation of the 2b-RAD method for genomic selection in scallop breeding. Scientific Reports 6(1), 19244. 10.1038/srep19244

Gaffney, P.M., Pascal, C.M., Barnhart, J., Grant, W.S., Seeb, J.E., 2010. Genetic homogeneity of weathervane scallops (*Patinopecten caurinus*) in the northeastern Pacific. Can. J. Fish. Aquat. Sci. 67(11), 1827–1839. 10.1139/f10-096

Gezan, S., de Oliveira, A.A., Galli, G., Murray, D., 2022. ASRgenomics: an R package with complementary genomic functions. VSN International, Hemel Hempstead, United Kingdom.

Gillespie, G.E., Bower, S.M., Marcus, K.L., Kieser, D., 2012. Biological synopsises for three exotic molluscs, Manila clam (*Venerupis philippinarum*), Pacific oyster (*Crassostrea gigas*) and Japanese scallop (*Mizuhopecten yessoensis*) licensed for aquaculture in British Columbia, in: DFO (Ed.). p. v + 97p.

Goddard, M.E., Hayes, B.J., 2009. Mapping genes for complex traits in domestic animals and their use in breeding programmes. Nature Reviews Genetics 10(6), 381–391. 10.1038/nrg2575

Gruber, B., Unmack, P.J., Berry, O.F., Georges, A., 2018. dartr: An R package to facilitate analysis of SNP data generated from reduced representation genome sequencing. Mol. Ecol. Resour. 18(3), 691–699. 10.1111/1755-0998.12745

Guo, X., 2009. Use and exchange of genetic resources in molluscan aquaculture. Reviews in Aquaculture 1(3-4), 251–259. 10.1111/j.1753-5131.2009.01014.x

Guo, X., Luo, Y., 2016. Scallops and scallop aquaculture in China, in: Shumway, S.E., Parsons, G.J. (Eds.), Scallops: Biology, ecology, aquaculture, and fisheries. Elsevier, pp. 937–952.

Guo, X., Puritz, J.B., Wang, Z., Proestou, D., Allen, S., Small, J., Verbyla, K., Zhao, H., Haggard, J., Chriss, N., Zeng, D., Lundgren, K., Allam, B., Bushek, D., Gomez-Chiarri, M., Hare, M., Hollenbeck, C., La Peyre, J., Liu, M., Lotterhos, K.E., Plough, L., Rawson, P., Rikard, S., Saillant, E., Varney, R., Wikfors, G., Wilbur, A., 2023. Development and evaluation of high-density SNP arrays for the eastern oyster *Crassostrea virginica*. Mar. Biotechnol. 25(1), 174–191. 10.1007/s10126-022-10191-3

Gutierrez, A.P., Matika, O., Bean, T.P., Houston, R.D., 2018. Genomic selection for growth traits in Pacific oyster (*Crassostrea gigas*): potential of low-density marker panels for breeding value prediction. Frontiers in Genetics 9. 10.3389/fgene.2018.00391

Gutierrez, A.P., Symonds, J., King, N., Steiner, K., Bean, T.P., Houston, R.D., 2020. Potential of genomic selection for improvement of resistance to ostreid herpesvirus in Pacific oyster (*Crassostrea gigas*). Anim. Genet. 51(2), 249–257. 10.1111/age.12909

Gutierrez, A.P., Turner, F., Gharbi, K., Talbot, R., Lowe, N.R., Peñaloza, C., McCullough, M., Prodöhl, P.A., Bean, T.P., Houston, R.D., 2017. Development of a medium density combined-species SNP array for Pacific and European oysters (*Crassostrea gigas* and *Ostrea edulis*). G3 Genes|Genomes|Genetics 7(7), 2209–2218. 10.1534/g3.117.041780

Hadfield, J.D., 2010. MCMC methods for multi-response generalized linear mixed models: the MCMCglmm R package. Journal of Statistical Software 33(2), 1–22. 10.18637/jss.v033.i02

Hartig, F., 2024. DHARMa: residual diagnostics for hierarchical (multi-level/ mixed) regression models., 0.4.7 ed.

Hedgecock, D., Davis, J.P., 2007. Heterosis for yield and crossbreeding of the Pacific oyster *Crassostrea gigas*. Aquaculture 272, S17–S29. 10.1016/j.aquaculture.2007.07.226

Hedgecock, D., Gaffney, P.M., Goulletquer, P., Guo, X., Reece, K., Warr, G.W., 2005. The case for sequencing the Pacific oyster genome. J. Shellfish Res. 24(2), 429–441, 413.

Hedgecock, D., Pudovkin, A.I., 2011. Sweepstakes reproductive success in highly fecund marine fish and shellfish: a review and commentary. Bull. Mar. Sci. 87(4), 971–1002.

Holden, J.J., Collicutt, B., Covernton, G., Cox, K.D., Lancaster, D., Dudas, S.E., Ban, N.C., Jacob, A.L., 2019. Synergies on the coast: challenges facing shellfish aquaculture development on the central and north coast of British Columbia. Mar. Policy 101, 108–117. 10.1016/j.marpol.2019.01.001

Hu, B., Tian, Y., Li, Q., Liu, S., 2022. Genomic signatures of artificial selection in the Pacific oyster, *Crassostrea gigas*. Evol. Appl. 15(4), 618–630. 10.1111/eva.13286

Jombart, T., Ahmed, I., 2011. adegenet 1.3-1: new tools for the analysis of genome-wide SNP data. Bioinformatics 27(21), 3070–3071. 10.1093/bioinformatics/btr521

Jourdan, A., Phocas, F., Boudry, P., Haffray, P., Allal, F., Maurouard, E., Heurtebise, S., Morga, B., Dégremont, L., Morvezen, R., 2026. Exploring genomic resistance to coinfection: single or dual pathogen infection by ostreid herpesvirus 1 and *Vibrio aestuarianus* in Pacific oysters *Crassostrea gigas*. Aquaculture, 743725. 10.1016/j.aquaculture.2026.743725

Kamaishi, T., Miwa, S., Goto, E., Matsuyama, T., Oseko, N., 2010. Mass mortality of giant abalone *Haliotis gigantea* caused by a *Francisella* sp. bacterium. Dis Aquat Organ 89(2), 145–154. 10.3354/dao02188

Kassambara, A., Kosinski, M., Biecek, P., 2024. survminer: drawing survival curves using ggplot2, 0.5.0 ed.

Kawahara, M., Kanamori, M., Meyer, G.R., Yoshinaga, T., Itoh, N., 2018. *Francisella halioticida*, identified as the most probable cause of adductor muscle lesions in Yesso scallops *Patinopecten yessoensis* cultured in southern Hokkaido, Japan. Fish Pathol. 53(2), 78–85. 10.3147/jsfp.53.78

Kawahara, M., Meyer, G.R., Lowe, G.J., Kim, E., Polinski, M.P., Yoshinaga, T., Itoh, N., 2019. Parallel studies confirm *Francisella halioticida* causes mortality in Yesso scallops *Patinopecten yessoensis*. Dis Aquat Organ 135(2), 127–134. 10.3354/dao03383

Knaus, B.J., Grünwald, N.J., 2017. vcfr: a package to manipulate and visualize variant call format data in R. Mol. Ecol. Resour. 17(1), 44–53. 10.1111/1755-0998.12549

Kosaka, Y., 2016. Scallop fisheries and aquaculture in Japan, in: Shumway, S.E., Parsons, G.J. (Eds.), Scallops: Biology, ecology, aquaculture, and fisheries. Elsevier, pp. 891–936.

Krzywinski, M., Schein, J., Birol, I., Connors, J., Gascoyne, R., Horsman, D., Jones, S.J., Marra, M.A., 2009. Circos: an information aesthetic for comparative genomics. Genome Res. 19(9), 1639–1645. 10.1101/gr.092759.109

Lenth, R.V., 2024. emmeans: estimated marginal means, aka least-squares means., 1.10.6 ed.

Li, H., 2013. Aligning sequence reads, clone sequences and assembly contigs with BWA-MEM. arXiv:1303.3997.

Liu, F., Li, Y., Yu, H., Zhang, L., Hu, J., Bao, Z., Wang, S., 2020. MolluscDB: an integrated functional and evolutionary genomics database for the hyper-diverse animal phylum Mollusca. Nucleic Acids Res. 49(D1), D988–D997. 10.1093/nar/gkaa918

Lou, R.N., Jacobs, A., Wilder, A.P., Therkildsen, N.O., 2021. A beginner’s guide to low-coverage whole genome sequencing for population genomics. Mol. Ecol. 30(23), 5966–5993. 10.1111/mec.16077

Meek, M.H., Larson, W.A., 2019. The future is now: Amplicon sequencing and sequence capture usher in the conservation genomics era. Mol. Ecol. Resour. 19(4), 795–803. 10.1111/1755-0998.12998

Meyer, G.R., Itoh, N., 2025. Infection with Francisella halioticida in Yesso scallops Mizuhopecten (=Patinopecten) yessoensis, in: Smolowitz, R. (Ed.), Diseases of Bivalves. Academic Press, pp. 65–70.

Meyer, G.R., Lowe, G.J., Gilmore, S.R., Bower, S.M., 2017. Disease and mortality among Yesso scallops *Patinopecten yessoensis* putatively caused by infection with *Francisella halioticida*. Dis. Aquat. Org. 125(1), 79–84.

Moran, B.M., Anderson, E.C., 2019. Bayesian inference from the conditional genetic stock identification model. Can. J. Fish. Aquat. Sci. 76(4), 551–560. 10.1139/cjfas-2018-0016

Normandeau, E., de Ronne, M., Torkamaneh, D., 2023. SNPLift: Fast and accurate conversion of genetic variant coordinates across genome assemblies. bioRxiv, 2023.2006.2013.544861. 10.1101/2023.06.13.544861

Paria, S.S., Rahman, S.R., Adhikari, K., 2022. fastman: A fast algorithm for visualizing GWAS results using Manhattan and Q-Q plots. bioRxiv, 2022.2004.2019.488738. 10.1101/2022.04.19.488738

Parsons, G.J., Lauzier, R.B., Bourne, N.F., 2016. Scallops of the West Coast of North America, in: Shumway, S.E., Parsons, G.J. (Eds.), Scallops: Biology, ecology, aquaculture, and fisheries. Elsevier, pp. 697–717.

Pernet, F., Lupo, C., Bacher, C., Whittington, R.J., 2016. Infectious diseases in oyster aquaculture require a new integrated approach. Philosophical Transactions of the Royal Society B: Biological Sciences 371(1689). 10.1098/rstb.2015.0213

Pew, J., Muir, P.H., Wang, J., Frasier, T.R., 2015. related: an R package for analysing pairwise relatedness from codominant molecular markers. Mol Ecol Resour 15(3), 557–561. 10.1111/1755-0998.12323

Plough, L.V., 2016. Genetic load in marine animals: a review. Current Zoology 62(6), 567–579. 10.1093/cz/zow096

Quayle, D.B., 1988. Pacific oyster culture in British Columbia. Department of Fisheries and Oceans, Ottawa.

Quinlan, A.R., Hall, I.M., 2010. BEDTools: a flexible suite of utilities for comparing genomic features. Bioinformatics 26(6), 841–842. 10.1093/bioinformatics/btq033

R Core Team, 2026. R: A Language and Environment for Statistical Computing. R Foundation for Statistical Computing, Vienna, Austria.

Robert, R., Gérard, A., 1999. Bivalve hatchery technology: The current situation for the Pacific oyster *Crassostrea gigas* and the scallop *Pecten maximus* in France. Aquat. Living Resour. 12(2), 121–130. 10.1016/S0990-7440(99)80021-7

Rochette, N.C., Rivera-Colón, A.G., Catchen, J.M., 2019. Stacks 2: Analytical methods for paired-end sequencing improve RADseq-based population genomics. Mol. Ecol. 28(21), 4737–4754. 10.1111/mec.15253

Saunders, R.G., Heath, W.A., 1994. New developments in scallop farming in British Columbia, Bull. Aquacult. Assoc. Can., 3 ed., pp. 3–7.

Sauvage, C., Bierne, N., Lapègue, S., Boudry, P., 2007. Single nucleotide polymorphisms and their relationship to codon usage bias in the Pacific oyster *Crassostrea gigas*. Gene 406(1), 13–22. 10.1016/j.gene.2007.05.011

Sutherland, B.J.G., Divilov, K., Thompson, N.F., Delomas, T.A., Lunda, S.L., Langdon, C.J., Green, T.J., 2026. Pedigree-based genome-wide imputation using a low-density amplicon panel for the highly polymorphic Pacific oyster *Crassostrea* (*Magallana*) *gigas*. Aquaculture 612, 743096. 10.1016/j.aquaculture.2025.743096

Sutherland, B.J.G., Itoh, N., Gilchrist, K., Boyle, B., Roth, M., Green, T.J., 2023. Genomic diversity of wild and cultured Yesso scallop *Mizuhopecten yessoensis* from Japan and Canada. G3 Genes|Genomes|Genetics 13(12), jkad242. 10.1093/g3journal/jkad242

Sutherland, B.J.G., Rycroft, C., Ferchaud, A.L., Saunders, R., Li, L., Liu, S., Chan, A.M., Otto, S.P., Suttle, C.A., Miller, K.M., 2020. Relative genomic impacts of translocation history, hatchery practices, and farm selection in Pacific oyster *Crassostrea gigas* throughout the Northern Hemisphere. Evol. Appl. 13(6), 1380–1399. 10.1111/eva.12965

Sutherland, B.J.G., Thompson, N.F., Surry, L.B., Reddy Gujjula, K., Carrasco, C.D., Chadaram, S., Lunda, S.L., Langdon, C.J., Chan, A.M., Suttle, C.A., Green, T.J., 2024. An amplicon panel for high-throughput and low-cost genotyping of Pacific oyster. G3: Genes|Genomes|Genetics 14(9), jkae125. 10.1093/g3journal/jkae125

Therneau, T.M., 2024. A package for survival analysis in R.

Therneau, T.M., Grambsch, P.M., 2000. Modeling survival data: extending the Cox model. Springer, New York.

Thompson, N.F., Sutherland, B.J.G., Green, T.J., Delomas, T.A., 2025. A free lunch: microhaplotype discovery in an existing amplicon panel improves parentage assignment for the highly polymorphic Pacific oyster. G3: Genes Genomes Genetics 15(2), jkae280. 10.1093/g3journal/jkae280

Vu, S.V., Gondro, C., Nguyen, N.T.H., Gilmour, A.R., Tearle, R., Knibb, W., Dove, M., Vu, I.V., Khuong, L.D., O’Connor, W., 2021. Prediction accuracies of genomic selection for nine commercially important traits in the Portuguese oyster (*Crassostrea angulata*) Using DArT-Seq technology. Genes 12(2). 10.3390/genes12020210

Wang, S., Zhang, J., Jiao, W., Li, J., Xun, X., Sun, Y., Guo, X., Huan, P., Dong, B., Zhang, L., Hu, X., Sun, X., Wang, J., Zhao, C., Wang, Y., Wang, D., Huang, X., Wang, R., Lv, J., Li, Y., Zhang, Z., Liu, B., Lu, W., Hui, Y., Liang, J., Zhou, Z., Hou, R., Li, X., Liu, Y., Li, H., Ning, X., Lin, Y., Zhao, L., Xing, Q., Dou, J., Li, Y., Mao, J., Guo, H., Dou, H., Li, T., Mu, C., Jiang, W., Fu, Q., Fu, X., Miao, Y., Liu, J., Yu, Q., Li, R., Liao, H., Li, X., Kong, Y., Jiang, Z., Chourrout, D., Li, R., Bao, Z., 2017. Scallop genome provides insights into evolution of bilaterian karyotype and development. Nature Ecology & Evolution 1(5), 0120. 10.1038/s41559-017-0120

Wang, Z., Casas, S., Peyre, J.L., Rikard, S., Williams, M.L., Tarnecki, A., Bushek, D., Guo, X., 2025. Genomic selection for Dermo resistance in the eastern oyster *Crassostrea virginica*: production and laboratory testing of F1 generation. J. Shellfish Res. 44(1), 75–87, 13.

Wickham, H., 2016. ggplot2: elegant graphics for data analysis. Springer-Verlag New York.

Xing, Q., Yang, Z., Zhu, X., Liu, J., Huang, X., Hu, J., Bao, Z., 2022. Interspecific hybridization between *Patinopecten yessoensis* (♀) and *P. caurinus* (♂) with heterosis in growth and temperature tolerance. Aquaculture 547, 737489. 10.1016/j.aquaculture.2021.737489

Yang, B., Li, Y., Li, Q., Liu, S., 2024a. High-throughput and cost-effective genotyping by low-coverage whole genome sequencing with genotype imputation in Pacific oyster, *Crassostrea gigas*. Aquaculture 591, 741134. 10.1016/j.aquaculture.2024.741134

Yang, B., Li, Y., Xu, M., Li, Q., Liu, S., 2026. Developing efficient and cost-effective NGS sequencing libraries for high-throughput genotyping by whole-genome resequencing in aquaculture. Aquaculture 613, 743431. 10.1016/j.aquaculture.2025.743431

Yang, B., Zhi, C., Li, P., Xu, C., Li, Q., Liu, S., 2024b. Genomic selection accelerates genetic improvement of resistance to vibriosis in the Pacific oyster, *Crassostrea gigas*. Aquaculture 584, 740679. 10.1016/j.aquaculture.2024.740679

Zhao, L., Li, Y., Li, Y., Yu, J., Liao, H., Wang, S., Lv, J., Liang, J., Huang, X., Bao, Z., 2017. A genome-wide association study identifies the genomic region associated with shell color in Yesso scallop, *Patinopecten yessoensis*. Mar. Biotechnol. 19, 301–309.

Zhou, X., Stephens, M., 2012. Genome-wide efficient mixed-model analysis for association studies. Nat. Genet. 44(7), 821–824. 10.1038/ng.2310

