## Supplemental Results for "An amplicon panel for high-throughput and low-cost genotyping of Yesso scallop *Mizuhopecten yessoensis*"

**Table of Contents:**


| | |
|---|---|
| Figure S1. Genotyping rate per sample (multiple datasets) | Page 2 |
| Figure S2. PCA for weathervane and Yesso scallops | Page 3 |
| Figure S3. DAPC for weathervane and Yesso scallops | Page 4 |
| Figure S4. Proportion of loci polymorphic in weathervane and Yesso scallops from BC and Japan | Page 5 |
| Figure S5. Per-family *F. halioticida* challenge survival probability | Page 6 |
| Figure S6. Per-family *F. halioticida* challenge survival (days post-exposure) and size | Page 7 |
| Figure S7. Model assumption plots for per-family size comparisons. | Page 8 |
| Table S1. Results of different marker selection methods | Page 9 |
| Table S2. Number of designed SNPs on each chromosome | Page 9 |
| Table S3. Results of SNP filtering steps | Page 10 |
| Table S4. Statistical testing of size differences between families | Page 11 |
| Table S5. Number of genotyped samples per family in *F. halioticida* trial | Page 12 |

**(A)**

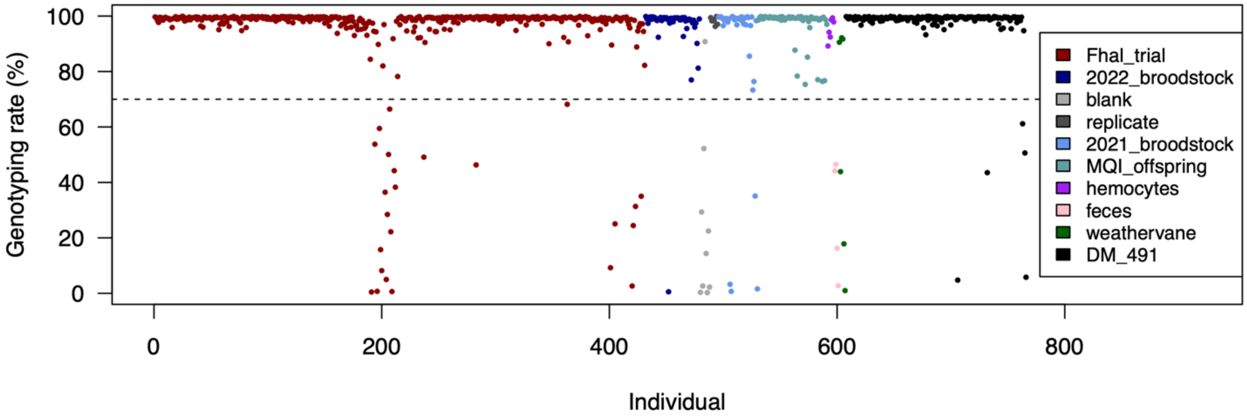


**(B)**

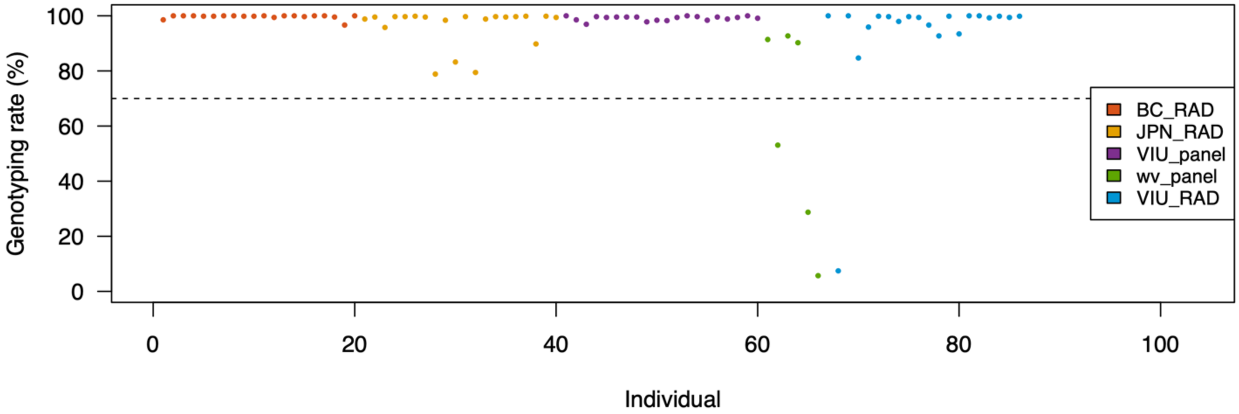


**Figure S1.** Genotyping rate (GR) per sample in (A) the pilot study using *de novo* SNP discovery; and (B) the analysis of weathervane scallop from Alaska and Yesso scallop from BC and Japan using ddRADseq or amplicon panel (*de novo*) data. In both datasets, samples below GR 70% were removed.

**(A)**

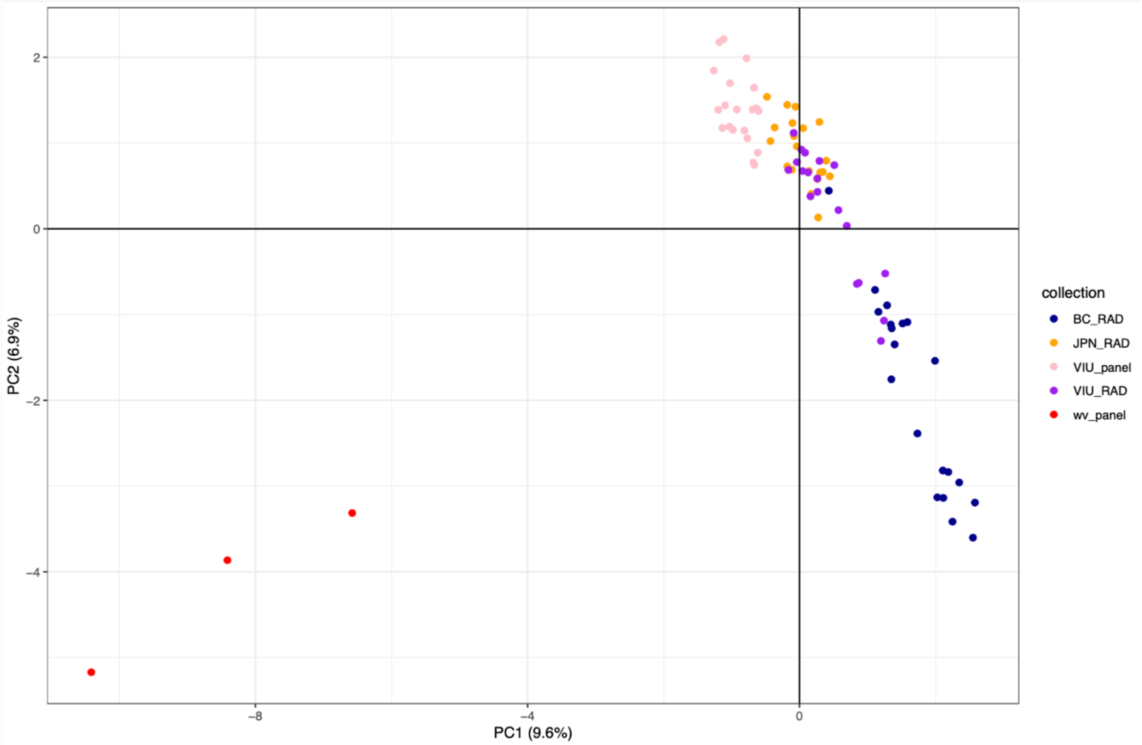


**(B)**

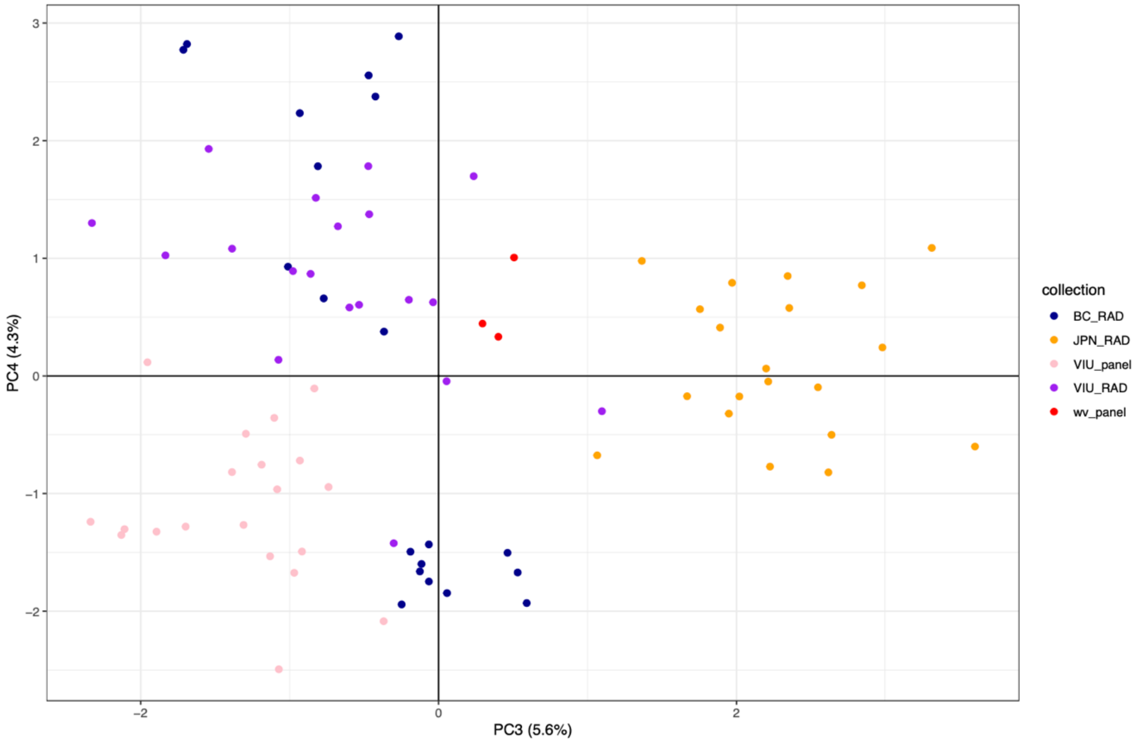


**Figure S2.** Principal components analysis (PCA) showing samples clustered by genotypes in PCs 1-4 in the weathervane vs. Yesso scallop analysis (RAD = ddRADseq; panel = amplicon panel).

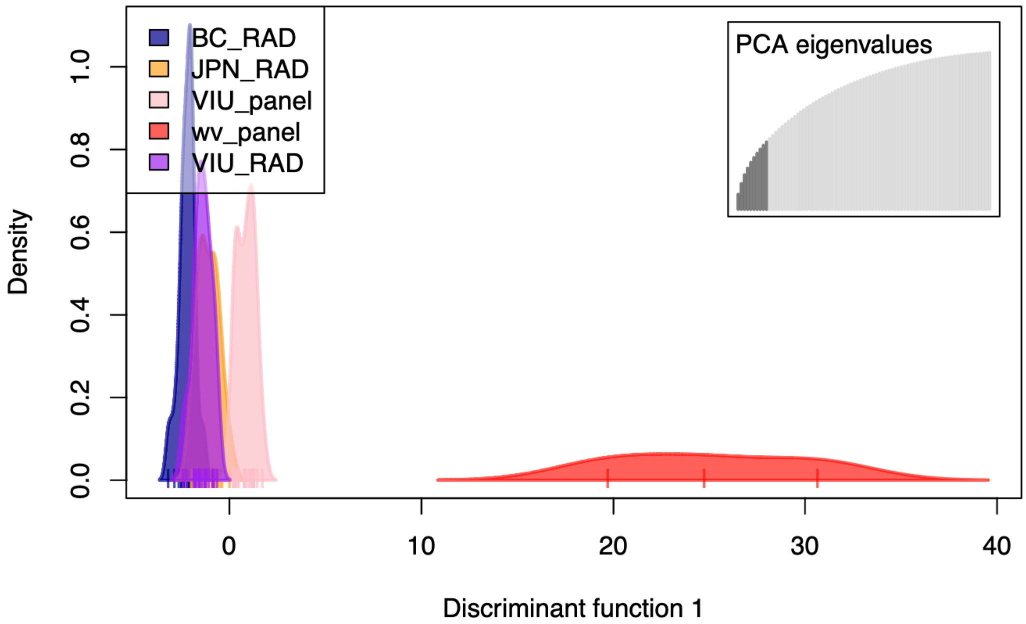


**Figure S3.** Supervised discriminant analysis of principal components (DAPC) conducted to separate populations in the Yesso scallop and weathervane scallop analysis. Yesso scallop samples from BC (cultivated) and Japan (wild) grouped near 0 on discriminant function 1, and the weathervane scallops grouped above 10 on DF1. The amplicon panel-based data for Yesso scallop grouped slightly separately from the reduced-representation dataset (RAD).

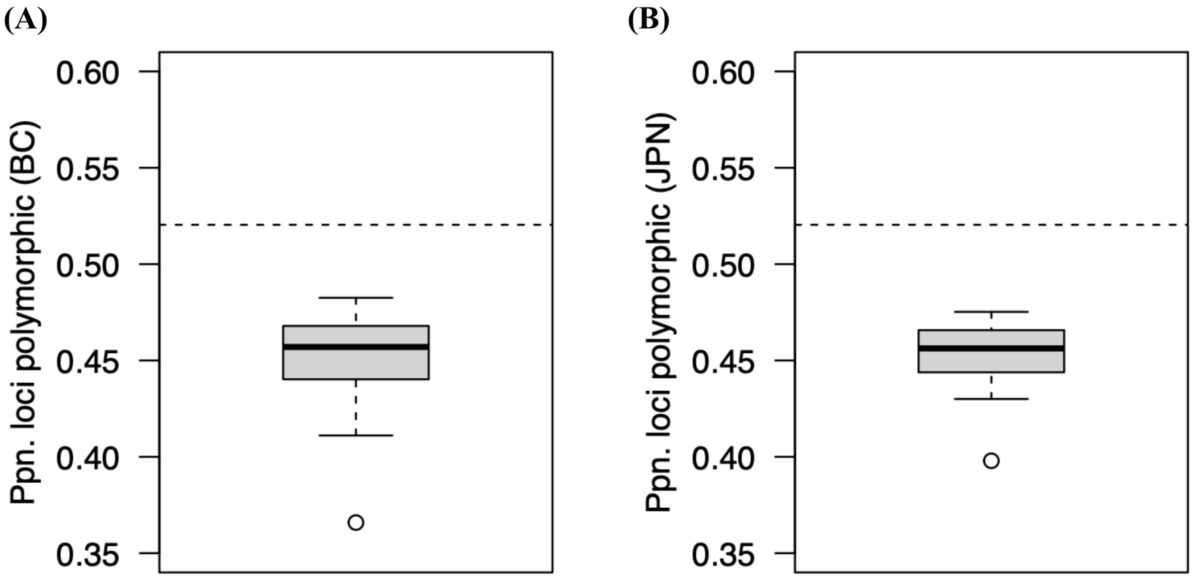


**Figure S4.** Proportion of polymorphic loci from the total genotyped loci (n = 686) in the Yesso scallop and weathervane scallop comparison study. The horizontal hatched line shows the proportion of polymorphic loci in the weathervane scallops (n = 3; 52.0%), and the distribution of values for 50 iterative subsamples of three individuals each subsample for (A) BC Yesso scallop samples; and (B) Japan Yesso scallop samples.

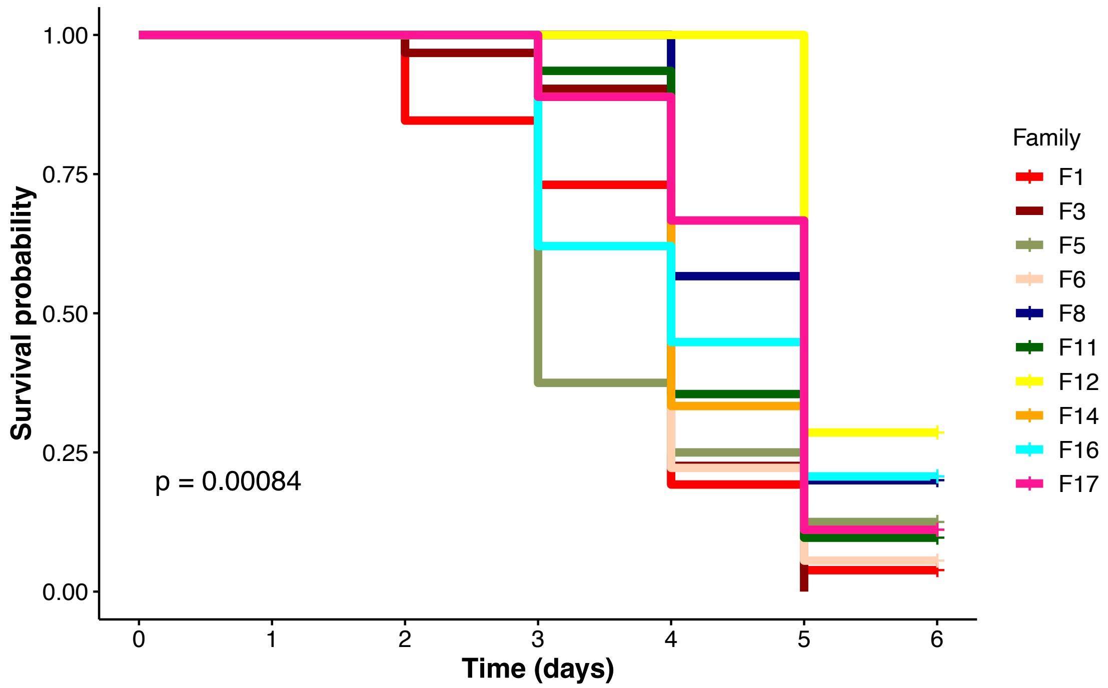


**Figure S5.** Kaplan-Meier survival probability over time (days) for each Yesso scallop family in response to injection with *Francisella halioticida* in the laboratory disease challenge. A value of Day 6 indicates survival through the entire trial. Families with n < 5 or with evidence of contamination from another family were not included in the analysis and are not shown here.

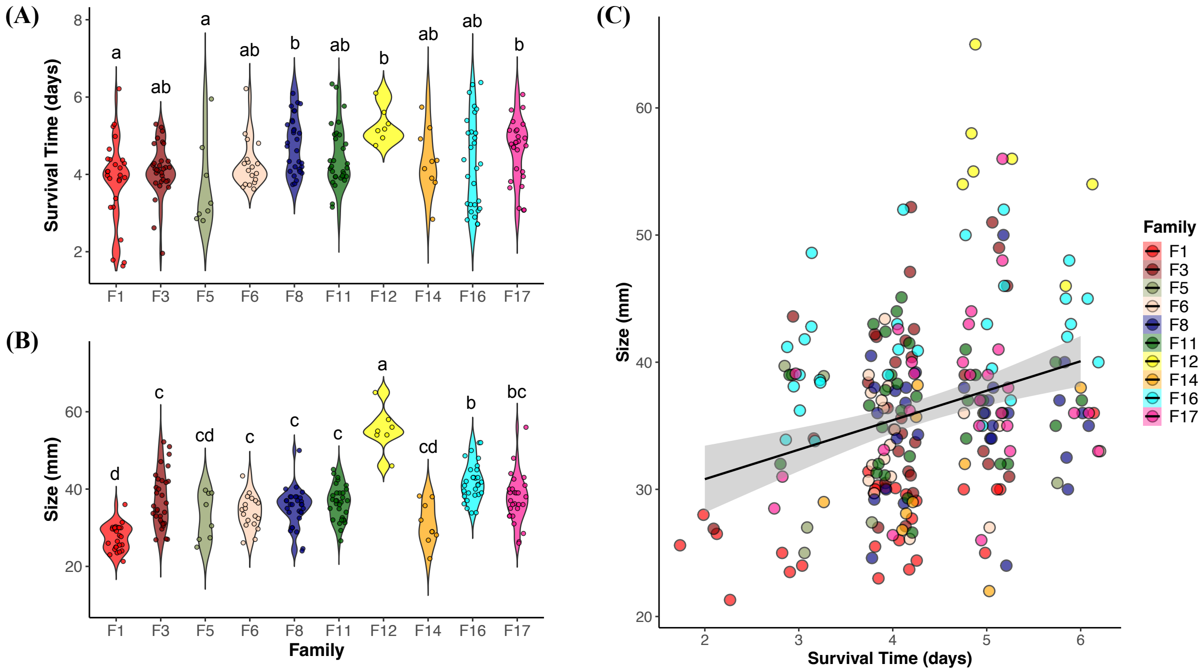


**Figure S6.** Violin plots of (A) survival (days post-exposure) of each scallop individual in each family in the *F. halioticida* challenge trial, along with (B) per-individual size (mm). (C) Survival showed a significant correlation with individual size in the infection trial. Families with n < 5 or with evidence of contamination from another family were not included in the analysis and are not shown here.

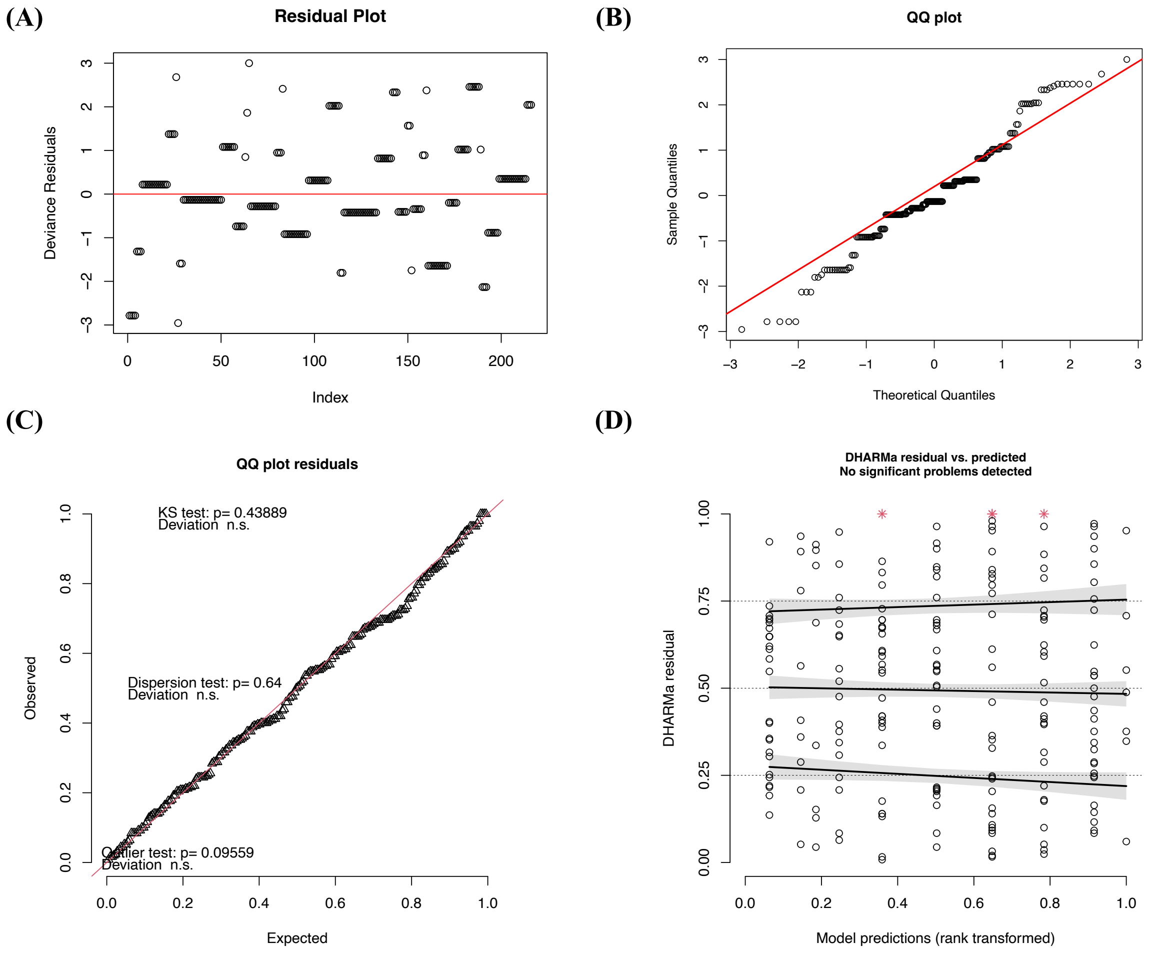


**Figure S7.** (A-B) Residual and Q-Q plot for parametric regression model for accelerated failure times using a log-logistic distribution of binary survival data. (C-D) DHARMa residual plots showing Q-Q plot and residuals compared with predicted values of a linear model of scallop size (mm) between families.

**Table S1.** Different marker selection methods varying number and size of windows.

| **Selection scenario** | Number markers selected | Average $H_{OBS}$ (VIU) | $H_{OBS}$ range (VIU) | Average $H_{OBS}$ (JPN) | $H_{OBS}$ range (JPN) |
|---|---|---|---|---|---|
| (i) 600 windows of 1.7 Mbp each | 569 | 0.34 | 0.03-0.49 | 0.33 | 0.03-0.49 |
| (ii) 200 windows of 5.0 Mbp each | 582 | 0.36 | 0.03-0.49 | 0.36 | 0.03-0.49 |
| (iii) 50 windows of 20.1 Mbp | 592 | 0.37 | 0.03-0.49 | 0.36 | 0.03-0.49 |

$H_{OBS}$ = observed heterozygosity

**Table S2.** Number of markers on each of the 20 chromosomes of the chromosome-level assembly.

| **Chromosome number** | **Chromosome name** | **Number of loci in design panel** |
|---|---|---|
| 1 | chr0 | 99 |
| 2 | chr1 | 35 |
| 3 | chr2 | 29 |
| 4 | chr3 | 31 |
| 5 | chr4 | 28 |
| 6 | chr5 | 34 |
| 7 | chr6 | 27 |
| 8 | chr7 | 28 |
| 9 | chr8 | 27 |
| 10 | chr9 | 29 |
| 11 | chr10 | 29 |
| 12 | chr11 | 25 |
| 13 | chr12 | 26 |
| 14 | chr13 | 23 |
| 15 | chr14 | 23 |
| 16 | chr15 | 22 |
| 17 | chr16 | 17 |
| 18 | chr17 | 22 |
| 19 | chr18 | 16 |
| 20 | chr19 | 19 |
| **TOTAL** | | **589** |

**Table S3.** Variant counts and percentages retained through the filtering process of *de novo* SNPs identified within amplicons. The full study dataset included all 768 barcoded samples, including nine blank samples. Filtering for minor allele frequency (MAF) was conducted within each of the pilot study subprojects. The Yesso vs. weathervane study used ddRADseq and panel data, including only a subset of the individuals (n = 86 total).

| | **Full Study** | | **Yesso vs. weathervane** | |
|---|---|---|---|---|
| **Filtering stage** | **No. variants** | **% retained** | **No. variants** | **% retained** |
| All variants | 2,983,897 | - | 445,934 | - |
| Retain variants > 5 bp from indel | 2,944,968 | 98.70% | 426,843 | 95.72% |
| Retain if missing in < 15%* of samples | 4,281 | 0.14% | 2,036 | 0.46% |
| Retain SNPs only | 3,630 | 0.12% | 1,702 | 0.24% |
| SNP quality ≥ 99 in at least one sample | 3,303 | 0.11% | 1,389 | 0.31% |
| Average depth across samples > 10 reads | 3,291 | 0.11% | 1,389 | 0.31% |
| Retain biallelic SNPs only | 3,242 | 0.11% | 1,347 | 0.30% |
| Per sample, per genotype set as missing if<br>depth <10 or >100K;<br>or if genotype quality < 20<br><br>Retain if missing in < 15%* of samples | 2,592 | 0.09% | 691 | 0.15% |

*cutoff applied for weathervane vs. Yesso = 10%

**Table S4.** Numbers of individuals genotyped for each family from the *F. halioticida* trial. The number genotyped is the number of individuals retained after filters were applied. Family F9 had samples from both challenge 3 (C3) and 4 (C4). Family F19 had samples in bucket 2 (B2) and bucket 8 (B8), for which the hatchery records were uncertain of reliability of expected parentage.

| **Family** | **Number genotyped** |
|---|---|
| F1 | 24 |
| F2 | 1 |
| F3 | 27 |
| F4 | 21 |
| F5 | 2 |
| F6 | 17 |
| F8 | 21 |
| F9 | 24 (C3), 117 (C4) |
| F11 | 26 |
| F12 | 4 |
| F13 | 1 |
| F14 | 2 |
| F15 | 2 |
| F16 | 25 |
| F17 | 25 |
| F18 | 17 |
| F19 | 5 (B2), 18 (B8) |
| F20 | 26 |
| TOTAL | 405 |
